# Non-neuronal expression of SARS-CoV-2 entry genes in the olfactory system suggests mechanisms underlying COVID-19-associated anosmia

**DOI:** 10.1101/2020.03.25.009084

**Authors:** David H. Brann, Tatsuya Tsukahara, Caleb Weinreb, Marcela Lipovsek, Koen Van den Berge, Boying Gong, Rebecca Chance, Iain C. Macaulay, Hsin-jung Chou, Russell Fletcher, Diya Das, Kelly Street, Hector Roux de Bezieux, Yoon-Gi Choi, Davide Risso, Sandrine Dudoit, Elizabeth Purdom, Jonathan S. Mill, Ralph Abi Hachem, Hiroaki Matsunami, Darren W. Logan, Bradley J. Goldstein, Matthew S. Grubb, John Ngai, Sandeep Robert Datta

**Author notes:** These authors contributed equally to this work (David H. Brann, Tatsuya Tsukahara, Caleb Weinreb).

## Abstract

Altered olfactory function is a common symptom of COVID-19, but its etiology is unknown. A key question is whether SARS-CoV-2 (CoV-2) – the causal agent in COVID-19 – affects olfaction directly by infecting olfactory sensory neurons or their targets in the olfactory bulb, or indirectly, through perturbation of supporting cells. Here we identify cell types in the olfactory epithelium and olfactory bulb that express SARS-CoV-2 cell entry molecules. Bulk sequencing revealed that mouse, non-human primate and human olfactory mucosa expresses two key genes involved in CoV-2 entry, ACE2 and TMPRSS2. However, single cell sequencing and immunostaining demonstrated ACE2 expression in support cells, stem cells, and perivascular cells; in contrast, neurons in both the olfactory epithelium and bulb did not express ACE2 message or protein. These findings suggest that CoV-2 infection of non-neuronal cell types leads to anosmia and related disturbances in odor perception in COVID-19 patients.

## Introduction

SARS-CoV-2 (CoV-2) is a pandemic coronavirus that causes the COVID-19 syndrome, which can include upper respiratory infection (URI) symptoms, severe respiratory distress, acute cardiac injury and death (*1–4*). CoV-2 is closely related to other beta-coronaviruses, including the causal agents in pandemic SARS and MERS (SARS-CoV and MERS-CoV, respectively) and endemic viruses typically associated with mild URI syndromes (hCoV-OC43 and hCoV-229E) (*5–7*). Clinical reports suggest that infection with CoV-2 is associated with high rates of disturbances in smell and taste perception, including anosmia (*8–12*). While many viruses (including coronaviruses) induce transient changes in odor perception due to inflammatory responses, in at least some cases COVID-related anosmia has been reported to occur in the absence of significant nasal inflammation or coryzal symptoms (*11, 13–15*). This observation suggests that CoV-2 might directly target odor processing mechanisms, although the specific means through which CoV-2 alters odor perception remains unknown.

CoV-2 — like SARS-CoV — infects cells through interactions between its spike (S) protein and the ACE2 protein on target cells. This interaction requires cleavage of the S protein, likely by the cell surface protease TMPRSS2, although other proteases (such as Cathepsin B and L, CTSB/CTSL) may also be involved (*4–6, 16–20*). Other coronaviruses use different cell surface receptors and proteases to facilitate cellular entry, including DPP4, FURIN and HSPA5 for MERS-CoV, ANPEP for HCoV-229E, TMPRSS11D for SARS-CoV (in addition to ACE2 and TMPRSS2), and ST6GAL1 and ST3GAL4 for HCoV-OC43 and HCoV-HKU1 (*6, 21–23*).

We hypothesized that identifying the specific olfactory cell types susceptible to direct CoV-2 infection (due to e.g., ACE2 and TMPRSS2 expression) would provide insight into possible mechanisms through which COVID-19 causes altered smell perception. The nasal epithelium is divided into a respiratory epithelium (RE) and olfactory epithelium (OE), whose functions and cell types differ. The nasal RE is continuous with the epithelium that lines much of the respiratory tract and is thought to humidify air as it enters the nose; main cell types include basal cells, ciliated cells, secretory cells (including goblet cells), and brush/microvillar cells (*24, 25*) (Figure 1). The OE, in contrast, is responsible for odor detection, as it houses mature olfactory sensory neurons (OSNs) that interact with odors via receptors localized on their dendritic cilia. OSNs are supported by sustentacular cells, which act to structurally support sensory neurons, phagocytose and/or detoxify potentially damaging agents, and maintain local salt and water balance (*26–28*); microvillar cells and mucus-secreting Bowman’s gland cells also play important roles in maintaining OE homeostasis and function (*24, 29*)(Figure 1). In addition, the OE contains globose basal cells (GBCs), which are primarily responsible for regenerating OSNs during normal epithelial turnover, and horizontal basal cells (HBCs), which act as reserve stem cells activated upon tissue damage (*30–32*). OSNs elaborate axons that puncture the cribriform plate at the base of the skull and terminate in the olfactory bulb, whose local circuits process olfactory information before sending it to higher brain centers (Figure 1).

**Fig. 1.**
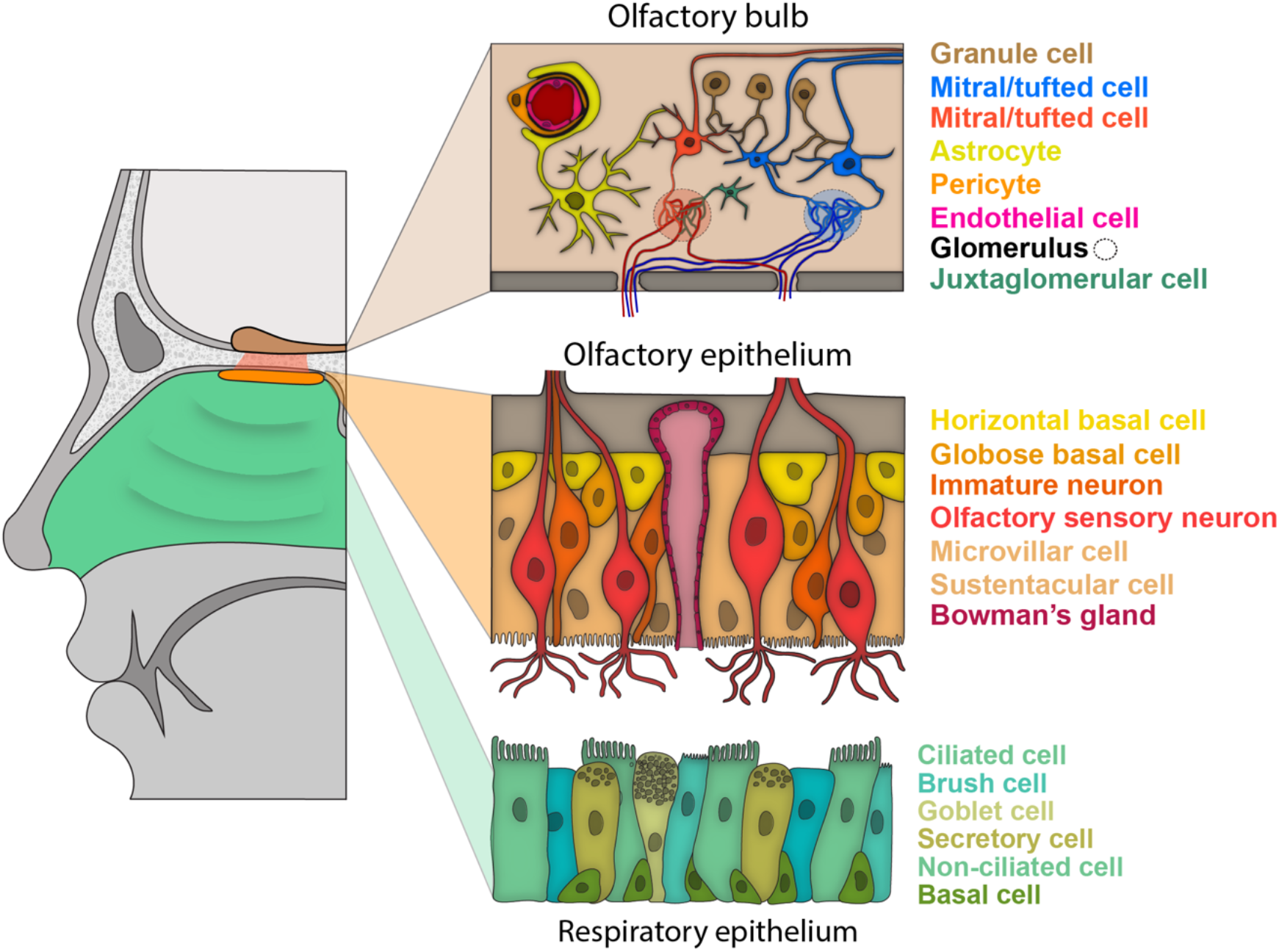
Schematic of the nasal respiratory and olfactory epithelium, and the olfactory bulb. Schematic of a sagittal view of the human nasal cavity, in which respiratory and olfactory epithelium are colored (left). For each type of epithelium, a schematic of the anatomy and known major cell types are shown (right). In the olfactory bulb in the brain (tan) the axons from olfactory sensory neurons coalesce into glomeruli, and mitral/tufted cells innervate these glomeruli and send olfactory projections to downstream olfactory areas. Glomeruli are also innervated by juxtaglomerular cells, a subset of which are dopaminergic.

It has recently been demonstrated through single cell RNA sequencing analysis (referred to herein as scSeq) that cells from the human upper airway — including nasal RE goblet, basal and ciliated cells—express high levels of ACE2 and TMPRSS2,suggesting that these RE cell types may serve as a viral reservoir during CoV-2 infection (*33*). However, analyzed samples in that dataset did not include any OSNs or sustentacular cells, indicating that tissue sampling in these experiments did not include the OE (*34, 35*). Here we query both new and previously published bulk RNA-Seq and scSeq datasets from the olfactory system for expression of ACE2, TMRPSS2 and other genes implicated in coronavirus entry. We find that non-neuronal cells in the OE and olfactory bulb, including support, stem and perivascular cells, express CoV-2 entry-associated transcripts and their associated proteins, suggesting that infection of these non-neuronal cell types contributes to anosmia in COVID-19 patients.

## Results

### Expression of CoV-2 entry genes in different cell types of the human olfactory epithelium

To determine whether genes relevant to CoV-2 entry are expressed in OSNs or other cell types in the human OE, we queried previously published bulk RNA-Seq data derived from the whole olfactory mucosa (WOM) of macaque, marmoset and human (*36*), and found expression of almost all CoV-entry-related genes in all WOM samples (Figure S1A). To identify the specific cell types in human OE that express ACE2, we quantified gene expression in scSeq derived from four human nasal biopsy samples recently reported by Durante et al (*37*). Neither ACE2 nor TMPRSS2 were detected in mature OSNs, whereas these genes were detected in both sustentacular cells and HBCs (Figures 2A-E). In contrast, genes relevant to cell entry of other CoVs were expressed in OSNs, as well as in other OE cell types. We confirmed the expression of ACE2 proteins via immunostaining of human olfactory epithelium biopsy tissue, which revealed expression in sustentacular and basal cells, and an absence of ACE2 protein in OSNs (Figures 2F and S1E). Together, these results demonstrate that sustentacular and olfactory stem cells, but not mature OSNs, are potentially direct targets of CoV-2 in the human OE.

**Fig. 2.**
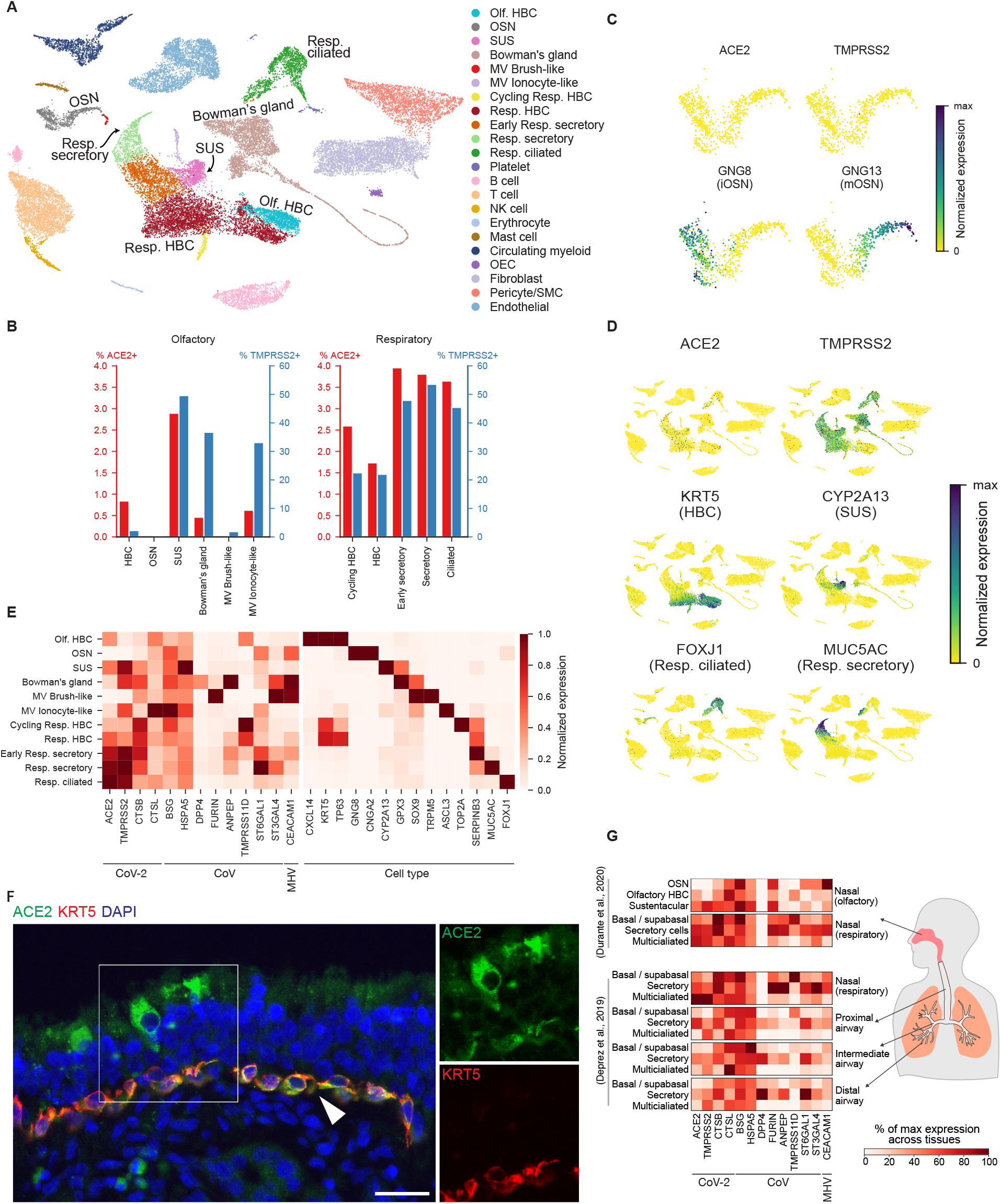
Coronavirus cell entry-related genes are expressed in human respiratory and olfactory epithelium but are not detected in human OSNs. **(A)** UMAP representation of cell types in human nasal biopsy scSeq data from Durante et al. 2020 (*37*). Each dot represents an individual cell, colored by cell type (HBC = horizontal basal cell, OSN = olfactory sensory neuron, SUS = sustentacular cell, MV: microvillar cell, Resp.: respiratory, OEC = olfactory ensheathing cell, SMC=smooth muscle cell). **(B)** Percent of cells expressing ACE2 and TMPRSS2. ACE2 was not detected in any OSNs, but was observed in SUS cells and HBCs, among other olfactory and respiratory epithelial cell types. Olfactory and respiratory cell types are shown separately. ACE2 and TMPRSS2 were also co-expressed above chance levels (Odds ratio 7.088, p-value 3.74E-57, Fisher’s exact test). **(C)** UMAP representations of 865 detected immature (GNG8) and mature (GNG13) OSNs. Neither ACE2 nor TMPRSS2 are detected in either population of OSNs. The color represents the normalized expression level for each gene (number of UMIs for a given gene divided by the total number of UMIs for each cell). **(D)** UMAP representations of all cells, depicting the normalized expression of CoV-2 related genes ACE2 and TMPRSS2, as well as several cell type markers. ACE2 and TMPRSS2 are expressed in respiratory and olfactory cell types, but not in OSNs. ACE2 and TMPRSS2 are detected in HBC (KRT5) and sustentacular (CYP2A13) cells, as well as other respiratory epithelial cell types, including respiratory ciliated (FOXJ1) cells. **(E)** Various CoV related genes including ACE2 and TMPRSS2, are expressed in respiratory and olfactory cell types, but not in OSNs. Gene expression for ACE2 and TMPRSS2 as well as marker genes for olfactory and respiratory epithelial cell types are shown normalized by their maximum expression across cell types. MHV, mouse hepatitis virus. **(F)** ACE2 immunostaining of human olfactory mucosal biopsy samples. ACE2 protein (green) is detected in sustentacular cells and KRT5-positive basal cells (red; white arrowhead). Nuclei were stained with DAPI (blue). Bar = 25 μm. The ACE2 and KRT5 channels from the box on the left are shown individually on the right **(G)** Gene expression across cell types and tissues in Durante et al. (top) and Deprez et al. (*34*)(bottom). Each gene is normalized to its maximum value across all tissues. Gene expression from Durante et al was normalized to that in Deprez et al to enable comparisons (see Methods and Figure S4). The tissues correspond to progressive positions along the airway from nasal to distal lung. ACE2 expression in olfactory HBC and sustentacular cells is comparable to that observed in other cell types in the respiratory tract.

Given that the nasopharynx is a major site of infection for CoV-2 (*10*), we compared the frequency of ACE2 and TMPRSS2 expression among the cell types in the human RE and OE (*37*). Sustentacular cells exhibited the highest frequency of ACE2 expression in the OE (2.90% of cells) although this frequency was slightly lower than that observed in respiratory ciliated and secretory cells (3.65% and 3.96%, respectively). While all HBC subtypes expressed ACE2, the frequency of expression of ACE2 was lower in olfactory HBCs (0.84% of cells) compared to respiratory HBCs (1.78% of cells) (Figure 2B). In addition, all other RE cell subtypes showed higher frequencies of ACE2 and TMPRSS2 expression than was apparent in OE cells.

These results demonstrate the presence of key CoV-2 entry-related genes in specific cell types in the OE, but at lower levels of expression than in RE isolated from the nasal mucosa. We wondered whether these lower levels of expression might nonetheless be sufficient for infection of CoV-2. It was recently reported that nasal RE has higher expression of CoV-2 entry genes than RE of the trachea or lungs (*38*), and we therefore asked where the OE fell within this previously established spectrum of expression. To address this question, we developed a two step alignment procedure in which we first sought to identify cell types that were common across the OE and RE, and then leveraged gene expression patterns in these common cell types to normalize gene expression levels across all cell types in the OE and RE (Figure S2). This approach revealed a correspondences between goblet cells in the RE and Bowman’s gland cells in the OE (96% mapping probability, see Methods), and between pulmonary ionocytes in the RE and a subset of microvillar cells in the OE (99% mapping probability, see Methods); after alignment, human OE sustentacular cells were found to express ACE2 and TMPRSS2 at levels similar to those observed in the remainder of the non-nasal respiratory tract (Figure 2G) (*38*). These results are consistent with the possibility that specific cell types in the human olfactory epithelium express ACE2 at a level that is permissive for direct infection.

### Expression of CoV-2 entry genes in different cell types of mouse olfactory epithelium

To further explore the distribution of CoV-2 cell entry genes in the olfactory system we turned to the mouse, which enables interrogative experiments not possible in humans. To evaluate whether that expression patterns observed in the mouse correspond to those observed in the human OE, we examined published datasets in which RNA-Seq was independently performed on mouse WOM and on purified populations of mature OSNs (*39–41*). The CoV-2 receptor Ace2 and the protease Tmprss2 were expressed in WOM, as were the cathepsins Ctsb and Ctsl (Figures 3A and S3A) (*39*). However, expression of these genes (with the exception of Ctsb) was much lower and Ace2 expression was nearly absent in purified OSN samples (Figures 3A and S3A, see Legend for counts). Genes used for cell entry by other CoVs (except St3gal4) were also expressed in WOM, and de-enriched in purified OSNs. The deenrichment of Ace2 and Tmprss2 in OSNs relative to WOM was also observed in two other mouse RNA-Seq datasets (*40, 41*) (Figure S3B). These data demonstrate that, as in humans, Ace2 and other CoV-2 entry-related genes are expressed in the mouse olfactory epithelium.

**Fig. 3.**
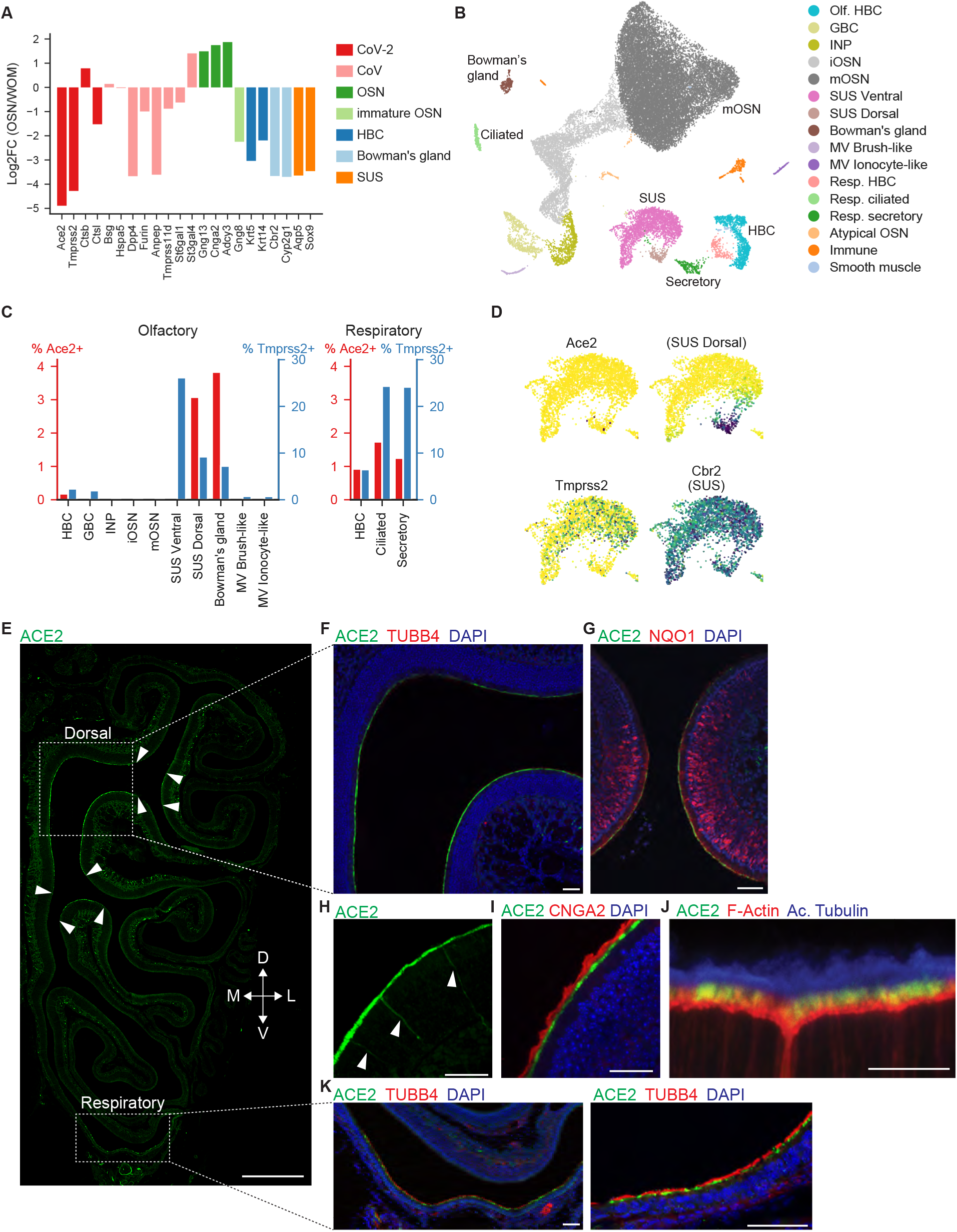
ACE2 is expressed in the mouse nasal epithelium but not in mature OSNs. **(A)** Log_2_-fold change (FC) in mean across-replicate gene expression between olfactory sensory neurons (OSNs) and whole olfactory mucosa (WOM) for coronavirus (CoV)-related genes and cell type markers (HBC = horizontal basal cells, SUS = sustentacular cells), data from Saraiva et al. (*39*). **(B)** UMAP representation of single cell transcriptome data from WOM, colored by cell types (mOSN: mature OSN, iOSN: immature OSN, INP: immediate neural precursor, GBC: globose basal cell, MV: microvillar cell, Resp.: respiratory). **(C)** Percent of cells expressing Ace2 and Tmprss2 in olfactory and respiratory cell types in the WOM (Drop-seq) dataset. Detection was considered positive if any transcripts (UMIs) were expressed for a given gene. Sustentacular cells (SUS) from dorsal and ventral zones are quantified separately. Ace2 is detected in dorsal sustentacular, Bowman’s gland, HBCs, as well as respiratory cell types. **(D)** UMAP representation of sustentacular cells, with expression of CoV-2 related genes Ace2 and Tmprss2, as well as marker genes for SUS (both pan-SUS marker Cbr2 and dorsal specific marker Sult1c1) indicated. Each point represents an individual sustentacular cell, and the color represents the normalized expression level for each gene (number of UMIs for a given gene divided by the total number of UMIs for each cell; in this plot Ace2 expression is binarized for visualization purposes). Ace2-positive sustentacular cells are found within the dorsal Sult1c1-positive subset. UMAP plots for other cell types are shown in Figure S2. **(E)** ACE2 immunostaining of mouse main olfactory epithelium. As shown in this epithelial hemisection, ACE2 protein is detected in the dorsal zone and respiratory epithelium. Note that the punctate Ace2 staining beneath the epithelial layer is likely associated with vasculature (see Figure 5F). Bar = 500 μm. Arrowheads depict the edges of ACE2 expression, corresponding to the presumptive dorsal zone (confirmed in **G**). Dashed boxes indicate the areas shown in **F** and **K** (left). **(F)** ACE2 protein is detected in the dorsal zone of the olfactory epithelium, which does not express the respiratory epithelial marker TUBB4. Bar = 50 μm. **(G)** Dorsal zone-specific expression of ACE2 in the olfactory epithelium was confirmed by co-staining with NQO1, a protein expressed in dorsal-zone OSNs. Bar = 50 μm **(H)** Bowman’s glands, which span from the lamina propria to the apical surface (arrowheads), were positive for ACE2 staining. Bar = 50 μm. **(I)** ACE2 signal in dorsal olfactory epithelium does not overlap with the cilia of olfactory sensory neurons, as visualized by CNGA2. Bar = 50 μm. **(J)** High magnification image of the apical end of the olfactory epithelium reveals that ACE2 signal is localized at the tip of villi of sustentacular cells, visualized by Phalloidin (F-Actin), but does not overlap with cilia of olfactory sensory neurons, as visualized by Acetylated Tubulin. Bar = 10 μm. **(K)** ACE2 expression in the respiratory epithelium was confirmed by co-staining with TUBB4. Bar = 50 μm.

The presence of Ace2 and Tmprss2 transcripts in mouse WOM and their (near total) absence in purified OSNs suggest that the molecular components that enable CoV-2 entry into cells are expressed in non-neuronal cell types in the mouse nasal epithelium. To identify the specific cell types that express Ace2 and Tmprss2, we performed scSeq (via Drop-seq, see Methods) on mouse WOM (Figure 3B). These results were consistent with observations made in the human epithelium: Ace2 and Tmprss2 were expressed in a fraction of sustentacular and Bowman’s gland cells, and a very small fraction of stem cells, but not in OSNs (zero of 17,666 identified mature OSNs, Figures 3C and S3C-D). Of note, only dorsally-located sustentacular cells, which express the markers Sult1c1 and Acsm4, were positive for Ace2 (Figures 3D and S3D-E). Indeed, reanalysis of the ACE2+ subset of human sustentacular cells revealed that all positive cells expressed genetic markers associated with the dorsal epithelium (Figure S1D). An independent mouse scSeq data set (obtained using the 10x Chromium platform, see Methods)revealed that olfactory sensory neurons did not express Ace2 (2 of 28769 mature OSNs were positive for Ace2), while expression was observed in a fraction of Bowman’s gland cells and HBCs (Figure S4, see methods). Expression in sustentacular cells was not observed in this dataset, which included relatively few dorsal sustentacular cells (a possible consequence of the specific cell isolation procedure associated with the 10x platform, which distinguishes it from Drop-seq; compare Figures S4C and 3D).

Staining of the mouse WOM with anti-ACE2 antibodies confirmed that ACE2 protein is expressed in sustentacular cells and is specifically localized to the sustentacular cell microvilli (Figure 3E-K). ACE2+ sustentacular cells were identified exclusively within the dorsal subregion of the OE; critically, within that region many (and possibly all) sustentacular cells expressed ACE2 (Figure 3F-G). This observation is consistent with the possibility that ACE2 protein can be broadly expressed in cell populations that exhibit sparse expression when characterized by scSeq. Staining was also observed in Bowman’s gland cells but not in OSNs (Figure 3H-J). Taken together, these data demonstrate that Ace2 is expressed by sustentacular cells that specifically reside in the dorsal epithelium in both mouse and human.

### Expression of CoV-2 entry genes in injured mouse olfactory epithelium

Viral injury can lead to broad changes in OE physiology that are accompanied by recruitment of stem cell populations tasked with regenerating the epithelium (*13, 30*). To characterize the distribution of Ace2 expression under similar circumstances, we injured the OE by treating mice with methimazole (which specifically ablates OSNs), and then employed a previously established lineage tracing protocol to perform scSeq on HBCs and their descendants during subsequent regeneration (see Methods) (*32*). This analysis revealed that after injury Ace2 and Tmprss2 are expressed in subsets of sustentacular cells and HBCs, as well as in the activated HBCs that serve to regenerate the epithelium (Figures 4A-C and S5; note that activated HBCs express Ace2 at higher levels than resting HBCs). Analysis of the Ace2+ sustentacular cell population revealed expression of dorsal epithelial markers (Figure 4D). To validate these results, we re-analyzed a similar lineage tracing dataset in which identified HBCs and their progeny were subject to Smart-seq2-based deep sequencing, which is more sensitive than scSeq (*32*). In this dataset, Ace2 was detected in more than 0.7% of GBCs, nearly 2% of activated HBCs and nearly 3% of sustentacular cells but was not detected in OSNs (Figures S5B). Furthermore, larger percentages of HBCs, GBCs and sustentacular cells expressed Tmprss2. Immunostaining with anti-ACE2 antibodies confirmed that ACE2 protein was present in activated stem cells under these regeneration conditions (Figure 4E). These results demonstrate that activated stem cells recruited during injury express ACE2, and do so at higher levels than those in resting stem cells.

**Fig. 4.**
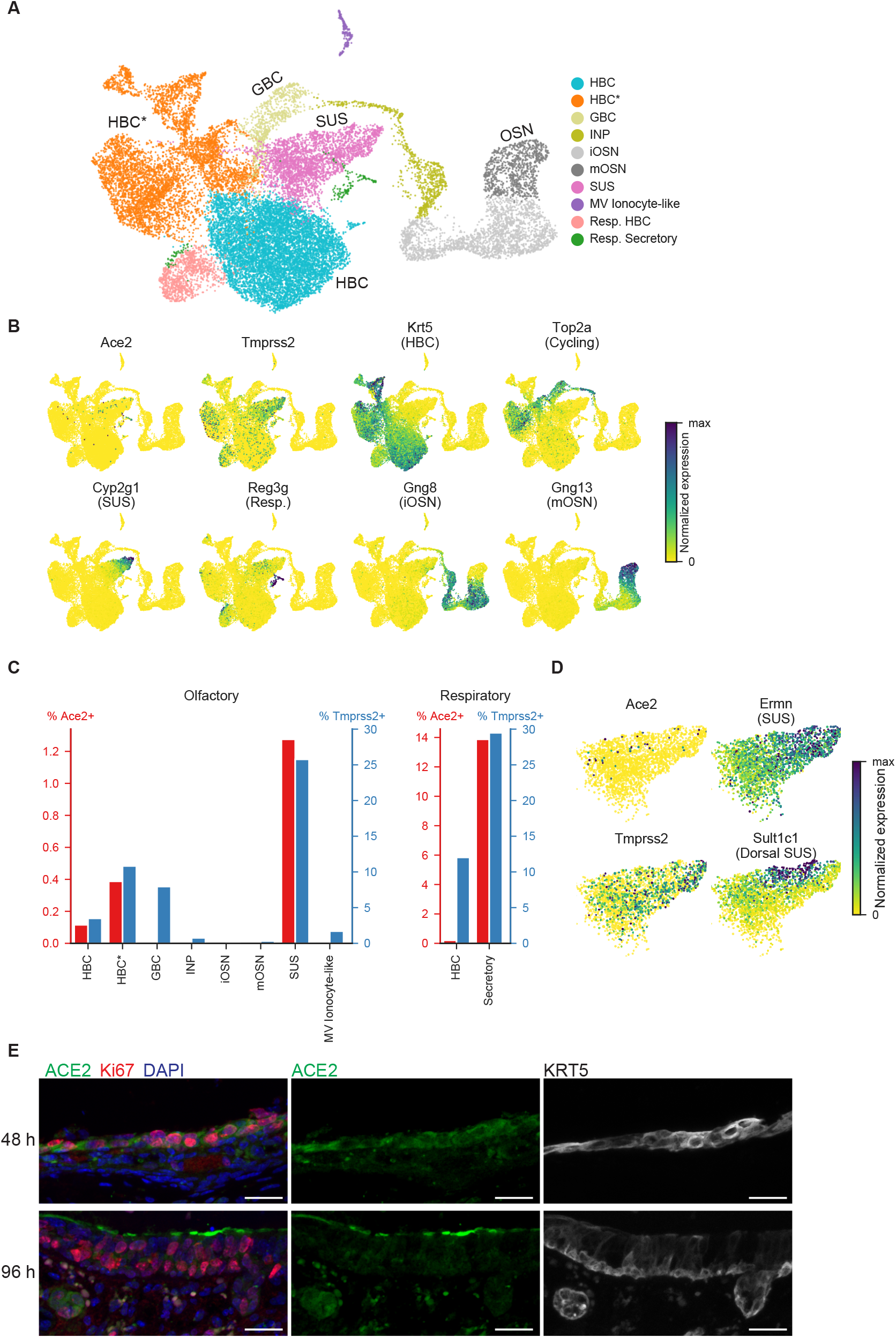
ACE2 is expressed in the mouse nasal epithelium in an injury model. **(A)** UMAP representation of single cell transcriptome data from an scSeq HBC lineage dataset, which includes several timepoints after epithelial injury induced by methimazole (mOSN: mature OSN, iOSN: immature OSN, INP: immediate neural precursor, SUS: sustentacular cell, GBC: globose basal cell, HBC: horizontal basal cell, HBC*: activated or cycling HBCs. MV: microvillar cell, Resp.: respiratory). **(B)** UMAP representation of the HBC lineage dataset, with cells expressing CoV-2 related genes Ace2 and Tmprss2, as well as marker genes for various cell types, indicated. The color represents normalized expression (number of UMIs for a given gene divided by the total number of UMIs for each cell). **(C)** Percent of cells expressing Ace2 and Tmprss2 in cell types identified in the HBC dataset. Ace2 is detected in sustentacular cells, HBC, activated/cycling HBC and respiratory cells. **(D)** UMAP representation of all sustentacular cells, indicating the normalized expression of CoV-2 related genes Ace2 and Tmprss2, as well as sustentacular (Ermn) cell markers. Ace2-positive sustentacular cells are largely a subset of dorsal SUS cells, as identified via the expression of Sult1c1. Sult1c1-positive sustentacular cells have higher levels of Ace2 (p=1.87E-03, Mann-Whitney test) and Ace2-positive sustentacular cells have higher levels of Sult1c1 (p=8.06E-07, Mann-Whitney test). **(E)** ACE2 immunostaining of mouse nasal epithelium after Methimazole treatment, together with cycling cell marker Ki67 and HBC marker KRT5. At 48 hours after injection, ACE2 signal is detected in Ki67+/KRT5+ activated HBCs (top). At 96 hours after injection, ACE2 signal is observed at the apical surface of Ki67+ cells (bottom). Because some of those cells still express low level of the HBC marker KRT5 and their staining pattern is reminiscent of dorsal sustentacular cells, these cells are likely early sustentacular cells in the process of differentiating from their stem cell precursors. Bar = 25 μm.

### Expression of CoV-2 entry genes in mouse olfactory bulb

Given the potential for the RE and OE in the nasal cavity to be directly infected with CoV-2, we assessed the expression of Ace2 and other CoV entry genes in the mouse olfactory bulb (OB), which is directly connected to OSNs via cranial nerve I (CN I); in principle, alterations in bulb function could cause anosmia independently of functional changes in the OE. To do so, we performed scSeq (using Drop-seq, see Methods) on the mouse OB, and merged these data with a previously published OB scSeq analysis, yielding a dataset with nearly 50,000 single cells (see Methods) (*42*). This analysis revealed that Ace2 expression was absent from OB neurons and instead was observed only in vascular cells, predominantly in pericytes, which are involved in blood pressure regulation, maintenance of the blood-brain barrier, and inflammatory responses (Figures 5A-D and S6–7) (*43*). Although other potential CoV proteases were expressed in the OB, Tmprss2 was not expressed.

**Fig. 5.**
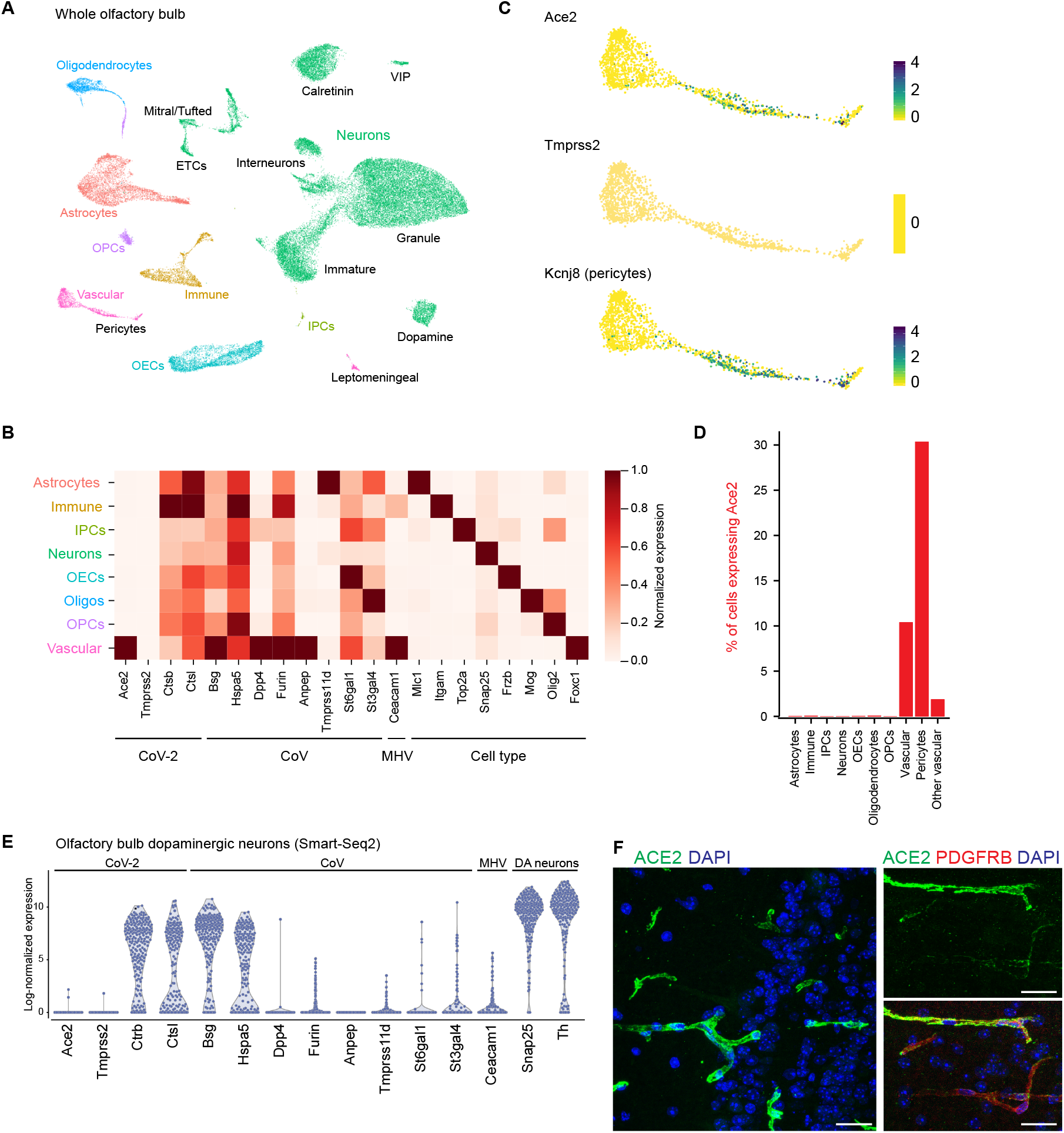
Expression of coronavirus entry genes in mouse olfactory bulb. **(A)** UMAP visualization of OB scSeq highlighting the main cell classes and subtypes, as observed in two integrated scSeq datasets (see Methods). VIP, vasoactive intestinal peptide positive neurons; ETCs, external tufted cells; OPCs, oligodendrocyte precursor cells; IPCs, intermediate precursor cells; OECs, olfactory ensheathing cells. Cluster information is summarized in Figures S6–7. **(B)** Heatmap showing expression of cell class markers and genes coding for coronavirus entry proteins in mouse olfactory bulb. Color scale shows scaled mean expression level per class, normalized by their maximum expression across cell types. Ace2 is specifically expressed in vascular cells. **(C)** UMAP representation of the vascular cell cluster showing expression of CoV-2 entry genes (Ace2, left; Tmprss2, center) and Kcnj8, a pericyte marker. The color scale depicts log-normalized UMI counts. **(D)** Percent of cells expressing ACE2. “Other vascular” denotes all vascular cells excluding pericytes. Ace2 expression is only detected in vascular cell types. **(E)** Violin plots showing Log_2_-normalized expression (Log_2_(TPM+1)) of coronavirus entry genes and dopaminergic neuron markers in manually sorted and deeply-sequenced single olfactory bulb dopaminergic neurons. Ace2 expression is not detected, suggesting that the lack of Ace2 expression in these cells in the Drop-seq and 10x datasets is not an artifact of undersampling. **(F)** ACE2 immunostaining of mouse main olfactory bulb confirms that ACE2 protein is present in vascular mural cells (left), including PDGFRB-positive pericytes (right), but is absent in neurons. Bar = 25 μm.

We also performed Smart-seq2-based deep sequencing of single OB dopaminergic juxtaglomerular neurons, a population of local interneurons in the OB glomerular layer that (like tufted cells) can receive direct monosynaptic input from nose OSNs (Figures 5E and S8, see Methods); these experiments confirmed the absence of Ace2 and Tmprss2 expression in this cell type. Immunostaining in the OB revealed that blood vessels expressed high levels of ACE2 protein, particularly in pericytes; consistent with the scSeq results, staining was not observed in any neuronal cell type (Figure 5F). These observations may also hold true for other brain regions, as re-analysis of 10 deeply sequenced scSeq datasets from different regions of the nervous system demonstrated that Ace2 and Tmprss2 expression is absent from neurons, consistent with prior immunostaining results (Figure S9)(*44*). Given the extensive similarities detailed above in expression patterns for Ace2 and Tmprss2 in the mouse and human, these findings (performed in mouse) suggest that OB neurons are likely not a primary site of infection, but that vascular pericytes may be sensitive to CoV-2 infection in the OB.

## Discussion

Here we show that subsets of OE sustentacular cells, HBCs, and Bowman’s gland cells in both mouse and human samples express the CoV-2 receptor ACE2 and the spike protein protease TMPRSS2. Human OE sustentacular cells express these genes at levels comparable to those observed in lung cells. In contrast, we failed to detect ACE2 expression in mature OSNs at either the transcript or protein levels. These observations suggest that CoV-2 does not directly enter OSNs, but instead may target OE support and stem cells. Similarly, neurons in the OB do not express ACE2, whereas vascular pericytes do. Thus primary infection of non-neuronal cell types — rather than sensory or bulb neurons — may be responsible for anosmia and related disturbances in odor perception in COVID-19 patients.

The identification of non-neuronal cell types in the OE and bulb susceptible to CoV-2 infection suggests four possible, non-mutually-exclusive mechanisms for the acute loss of smell reported in COVID-19 patients. First, local infection of support and vascular cells in the nose and bulb could cause significant inflammatory responses whose downstream effects could block effective odor conduction, or alter the function of OSNs or bulb neurons (*14*) (*45*). Second, damage to support cells (which are responsible for local water and ion balance) could indirectly influence signaling from OSNs to the brain (*46*). Third, damage to sustentacular cells and Bowman’s gland cells in mouse models can lead to diffuse architectural damage to the entire OE, which in turn could abrogate smell perception (*47*). Finally, vascular damage could lead to hypoperfusion and inflammation leading to changes in OB function.

Immunostaining in the mouse suggests that Ace2 protein is (nearly) ubiquitously expressed in sustentacular cells in the dorsal OE, despite sparse detection of Ace2 transcripts using scSeq. Similarly, nearly all vascular cells positive for a pericyte marker also expressed Ace2 protein, although only a fraction of OB pericytes were positive for Ace2 message when assessed using scSeq. Although Ace2 transcripts were more rarely detected than protein, there was a clear concordance at the cell type level: expression of Ace2 mRNA in a particular cell type accurately predicted the presence of Ace2 protein, while Ace2 transcript-negative cell types (including OSNs) did not express Ace2 protein. If humans also exhibit a similar relationship between mRNA and protein (a reasonable possibility given the precise match in olfactory cell types that express CoV-2 cell entry genes between the two species), then ACE2 protein is likely to be broadly expressed in human dorsal sustentacular cells. Thus, in the there may be many sustentacular cells available for CoV-2 infection in the human epithelium (which in turn could recruit a diffuse inflammatory process). That said, it remains possible that damage to the OE could be caused by more limited cell infection. For example, infection of subsets of sustentacular cells by the SDAV coronavirus in rats ultimately leads to disruption of the global architecture of the OE, suggesting that focal coronavirus infection may be sufficient to cause diffuse epithelial damage (*47*).

The natural history of CoV2-induced anosmia is only now being defined; while recovery of smell has been reported, it remains unclear whether in a subset of patients smell disturbances will be long-lasting or permanent (*8–12, 48*). We observe that activated HBCs, which are recruited after injury, express Ace2 at higher levels than those apparent in resting stem cells. While on its own it is likely that infection of stem cells would not cause acute smell deficits, in the context of infection the dual challenge of loss of sustentacular cells, together with the inability to effectively renew the OE over time, could result in persistent anosmia.

Many viruses, including coronaviruses, have been shown to propagate from the nasal epithelium past the cribriform plate to infect the OB; this form of central infection has been suggested to mediate olfactory deficits, even in the absence of lasting OE damage (*18, 49–53*). The rodent coronavirus MHV passes from the nose to the bulb, even though rodent OSNs do not express CEACAM1, the main MHV receptor (*50, 54*) (Figures S3C, S4E, S5A), suggesting that CoVs in the nasal mucosa can reach the brain through mechanisms independent of axonal transport by sensory nerves; interestingly, OB dopaminergic juxtaglomerular cells express CEACAM1 (Figure 4E), which likely supports the ability of MHV to target the bulb and change odor perception. One speculative possibility is that local seeding of the OE with CoV-2-infected cells can result in OSN-independent transfer of virions from the nose to the bulb, perhaps via the vascular supply shared between the OB and the OSN axons that comprise CN I. Although CN I was not directly queried in our datasets, it is reasonable to infer that vascular pericytes in CN I also express ACE2, which suggests a possible route of entry for CoV-2 from the nose into the brain. Given the absence of ACE2 in OB neurons, we speculate that any central olfactory dysfunction in COVID-19 is the secondary consequence of pericyte-mediated vascular inflammation (*43*).

We note several caveats that temper our conclusions. Although current data suggest that ACE2 is the most likely receptor for CoV-2 in vivo, it is possible (although it has not yet been demonstrated) that other molecules such as BSG may enable CoV-2 entry independently of ACE2 (Figures 2E, S3C, S4E, S5A) (*55, 56*). In addition, it has recently been reported that low level expression of ACE2 can support CoV-2 cell entry (*57*); it is possible, therefore, that ACE2 expression beneath the level of detection in our assays may yet enable CoV-2 infection of apparently ACE2 negative cell types. We also propose that damage to the olfactory system is either due to primary infection or secondary inflammation; it is possible (although has not yet been demonstrated) that cells infected with CoV-2 can form syncytia with cells that do not express ACE2. Such a mechanism could damage neurons adjacent to infected cells.

Any reasonable pathophysiological mechanism for COVID-19-associated anosmia must account for the high penetrance of smell disorders relative to endemic viruses, the apparent suddenness of smell loss (which can precede the development of other symptoms), and the transient nature of dysfunction in many patients (*8–12*) (*11, 13–15*); definitive identification of the disease mechanisms underlying COVID-19-mediated anosmia will require additional research. Nonetheless, our identification of cells in the OE and OB expressing molecules known to be involved in CoV-2 entry illuminates a path forward for future studies.

## Materials and Methods

### Human nasal scSeq dataset

Human scSeq data from Durante et al. (*37*) was downloaded from the GEO at accession GSE139522. 10x Genomics mtx files were filtered to remove any cells with fewer than 500 total counts. Additional preprocessing was performed as described above, including total counts normalization and filtering for highly variable genes using the SPRING gene filtering function “filter_genes” with parameters (90, 3, 10). The resulting data were visualized in SPRING and partitioned using Louvain clustering on the SPRING k-nearest-neighbor graph. Four clusters were removed for quality control, including two with low total counts (likely background) and two with high mitochondrial counts (likely stressed or dying cells). Putative doublets were also identified using Scrublet and removed (7% of cells). The remaining cells were projected to 40 dimensions using PCA. PCA-batch-correction was performed using Patient 4 as a reference, as previously described (*58*). The filtered data were then re-partitioned using Louvain clustering on the SPRING graph and each cluster was annotated using known marker genes, as described in (*37*). For example, immature and mature OSNs were identified via their expression of GNG8 and GNG13, respectively. HBCs were identified via the expression of KRT5 and TP63 and olfactory HBCs were distinguished from respiratory HBCs via the expression of CXCL14 and MEG3. Identification of SUS cells (CYP2A13, CYP2J2), Bowman’s gland (SOX9, GPX3), and MV ionocytes-like cells (ASCL3, CFTR, FOXI1) was also performed using known marker genes. For visualization, the top 40 principal components were reduced to two dimensions using UMAP with parameters (n_neighbors=15, min_dist=0.4).

The filtered human scSeq dataset contained 33358 cells. Each of the samples contained cells from both the olfactory and respiratory epithelium, although the frequency of OSNs and respiratory cells varied across patients, as previously described (*37*). 295 cells expressed ACE2 and 4953 cells expressed TMPRSS2. Of the 865 identified OSNs, including both immature and mature cells, none of the cells express ACE2 and only 2 (0.23%) expressed TMPRSS2. In contrast, ACE2 was reliably detected in at least 2% and TMPRSS2 was expressed in close to 50% of multiple respiratory epithelial subtypes. The expression of both known cell type markers and known CoV-related genes was also examined across respiratory and olfactory epithelial cell types. For these gene sets, the mean expression in each cell type was calculated and normalized by the maximum across cell types.

### Mapping scSeq datasets to each other

Data from Deprez et al. (*34*) were downloaded from the Human Cell Atlas website (https://www.genomique.eu/cellbrowser/HCA/; “Single-cell atlas of the airway epithelium (Grch38 human genome)”). A subset of these data was combined with a subset of the Durante data for mapping between cell types. For the Deprez data, the subset consisted of samples from the nasal RE that belonged to a cell type with >20 cells, including Basal, Cycling Basal, Suprabasal, Secretory, Mucous Multiciliated cells, Multiciliated, SMS Goblet and Ionocyte. We observed two distinct subpopulations of Basal cells, with one of the two populations distinguished by expression of Cxcl14. The cells in this population were manually identified using SPRING and defined for downstream analysis as a separate cell type annotation called “Basal (Cxcl14+)”. For the Durante data, the subset consisted of cells from cell types that had some putative similarity to cells in the Deprez dataset, including Olfactory HBC, Cycling respiratory HBC, Respiratory HBC, Early respiratory secretory cells, Respiratory secretory cells, Sustentacular cells, Bowman’s gland, Olfactory microvillar cells.

To establish a cell type mapping:

1. Durante (*37*) and Deprez (*34*) data were combined and gene expression values were linearly scaled so that all cells across datasets had the same total counts. PCA was then performed using highly variable genes (n=1477 genes) and PCA-batch-correction (*58*) with the Durante data as a reference set.
2. Mapping was then performed bidirectionally between the two datasets. Each cell from “Dataset 1” ‘voted’ for the 5 most similar cells in the “Dataset 2”, using distance in PCA space as the measure of similarity. A table *T* counting votes across cell types was then computed, where for cell type *i* in the Dataset 1 and cell type *j* in the Dataset 2,

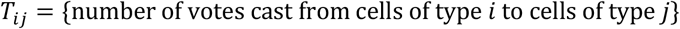 Thus, if Dataset 1 has *N* cells, then *T* would count 5*N votes (∑*T_ij_* = 5*N*)
3. The table of votes *T* was Z-scored against a null distribution, generated by repeating the procedure above 1000 times with shuffled cell type labels.

The resulting Z-scores were similar between the two possible mapping directions (Durante -> Deprez vs. Deprez -> Durante; R=0.87 Pearson correlation of mapping Z-scores). The mapping Z-scores were also highly robust upon varying the number of votes-cast per cell (R>0.98 correlation of mapping Z-scores upon changing the vote numbers to 1 or 50 as opposed to 5). Only cell-type correspondences with a high Z-score in both mapping directions (Z-score > 25) were used for downstream analysis.

To establish a common scale of gene expression between datasets, we restricted to cell type correspondences that were supported both by bioinformatic mapping and shared a nominal cell type designation based on marker genes. These included: Basal/suprabasal cells = “respiratory HBCs” from Durante et al., and “basal” and “suprabasal” cells from Deprez et al. Secretory cells = “early respiratory secretory cells” and “respiratory secretory cells” from Durante et al., and “secretory” cells from Deprez et al. Multiciliated cells = “respiratory ciliated cells” from Durante et al., and “multiciliated” cells from Deprez et al.

We next sought a transformation of the Durante data so that it would agree with the Deprez data within the corresponding cell types identified above To account for differing normalization strategies applied to each dataset prior to download (log normalization and rescaling with cell-specific factors for Deprez et al. but not for Durante et al.), we used the following ansatz for the transformation, where the pseudocount *p* is a global latent parameter and the rescaling factors *f_i_* are fit to each gene separately. In the equation below, *T* denotes the transformation and *e_ij_* represents a gene expression value for cell *i* and gene *j* in the Durante data:

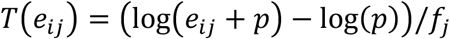

The parameter *p* was fit by maximizing the correlation of average gene expression across all genes between each of the cell type correspondences listed above. The rescaling factors *f_i_* were then fitted separately for each gene by taking the quotient of average gene expression between the Deprez data and the log-transformed Durante data, again across the cell type correspondences above.

### Mouse bulk RNA-Seq datasets

Normalized gene expression tables were obtained from previous published datasets (*36, 39–41*). For the mouse data sets, the means of the replicates from WOM or OSN were used to calculate Log_2_ fold changes. For the mouse data from Saraiva et al. and the primate data sets (*36, 39*), the normalized counts of the genes of interest from individual replicates were plotted. Below is a table with detailed sample information.

Sample information for the bulk RNA-seq data analyzed in this study

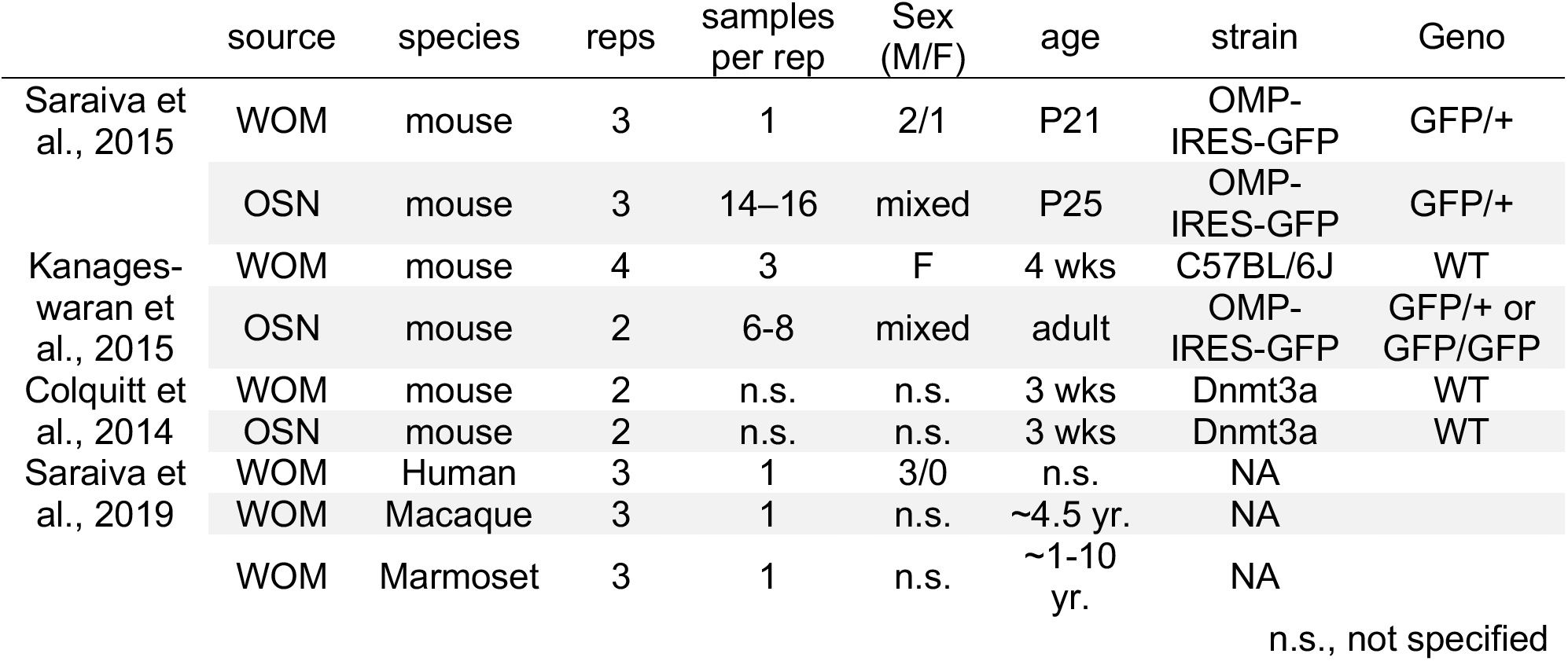

|  | source | species | reps | samples<br>per rep | Sex<br>(M/F) | age | strain | Geno |
| --- | --- | --- | --- | --- | --- | --- | --- | --- |
| Saraiva et al., 2015 | WOM | mouse | 3 | 1 | 2/1 | P21 | OMP-IRES-GFP | GFP/+ |
|  | OSN | mouse | 3 | 14–16 | mixed | P25 | OMP-IRES-GFP | GFP/+ |
| Kanageswaran et al., 2015 | WOM | mouse | 4 | 3 | F | 4 wks | C57BL/6J | WT |
|  | OSN | mouse | 2 | 6-8 | mixed | adult | OMP-IRES-GFP | GFP/+ or GFP/GFP |
| Colquitt et al., 2014 | WOM | mouse | 2 | n.s. | n.s. | 3 wks | Dnmt3a | WT |
|  | OSN | mouse | 2 | n.s. | n.s. | 3 wks | Dnmt3a | WT |
| Saraiva et al., 2019 | WOM | Human | 3 | 1 | 3/0 | n.s. | NA |  |
|  | WOM | Macaque | 3 | 1 | n.s. | ~4.5 yr. | NA |  |
|  | WOM | Marmoset | 3 | 1 | n.s. | ~1-10 yr. | NA |  |

n.s., not specified

### Mouse WOM Drop-seq experiments

#### Tissue dissection and single-cell dissociation for nasal epithelium

A new dataset of whole olfactory mucosa scSeq was generated from adult male mice (8–12 weeks-old). All mouse husbandry and experiments were performed following institutional and federal guidelines and approved by Harvard Medical School’s Institutional Animal Care and Use Committee (IACUC). Briefly, dissected main olfactory epithelium were cleaned up in 750 μl of EBSS (Worthington) and epithelium tissues were isolated in 750 μL of Papain (20 U/mL in EBSS) and 50 μL of DNase I (2000 U/mL). Tissue pieces were transferred to a 5 mL round-bottom tube (BD) and 1.75 mL of Papain and 450 μL of DNase I were added. After 1–1.5 hour incubation with rocking at 37°C, the suspension was triturated with a 5 mL pipette 15 times and passed through 40 μm cell strainer (BD) and strainer was washed with 1 mL of DMEM + 10 % FBS (Invitrogen). The cell suspension was centrifuged at 300g for 5 min. Cells were resuspended with 4 mL of DMEM + 10 % FBS and centrifuged at 300g for 5 min. Cells were suspended with PBS + 0.01 % BSA and concentration was measured by hemocytometer.

#### Drop-seq experiments

Drop-seq experiments were performed as previously described (*59*). Microfluidics devices were obtained from FlowJEM and barcode beads were obtained from chemgenes. 8 of 15 min Drop-seq runs were collected in total, which were obtained from 5 mice.

#### Sequencing of Drop-seq samples

8 replicates of Drop-seq samples were sequenced across 5 runs on an Illumina NextSeq 500 platform. Paired end reads from the fastq files were trimmed, aligned, and tagged via the Drop-seq tools (v1.13) pipeline, using STAR (v2.5.4a) with genomic indices from Ensembl Release 93. The digital gene expression matrix was generated for 4,000 cells for 0126_2, 5,000 cells for 0105, 0126_1, 051916_DS11, 051916_DS12, 051916_DS22, 5,500 cells for 051916_DS21, and 9,500 cells for 0106.

#### Preprocessing of Drop-seq samples

Processing of the WOM Drop-seq samples was performed in Seurat (v2.3.1). Cells with less than 500 UMIs or more than 15,000 UMIs, or higher than 5% mitochondrial genes were removed. Potential doublets were removed using Scrublet. Cells were initially preprocessed using the Seurat pipeline. Variable genes “FindVariableGenes” (y.cutoff = 0.6) were scaled (regressing out effects due to nUMI, the percent of mitochondrial genes, and replicate ids) and the data was clustered using 50 PCs with the Louvain algorithm (resolution=0.8). In a fraction of sustentacular cells, we observed co-expression of markers for sustentacular cells and other cell types (e.g. OSNs). Re-clustering of sustentacular cells alone separately out these presumed doublets from the rest of the sustentacular cells, and the presumed doublets were removed for the analyses described below.

#### Processing of Drop-seq samples

The filtered cells from the preprocessing steps were reanalyzed in python using Scanpy and SPRING. In brief, the raw gene counts in each cell were total counts normalized and variable genes were identified using the SPRING gene filtering function “filter_genes” with parameters (85, 3, 3); mitochondrial and olfactory receptor genes were excluded from the variable gene lists. The resulting 2083 variable genes were z-scored and the dimensionality of the data was reduced to 35 via principal component analysis. The k-nearest neighbor graph (n_neighbors=15) of these 35 PCs was clustered using the leiden algorithm (resolution=1.2) and was reduced to two dimensions for visualization via the UMAP method (min_dist=0.42). Clusters were manually annotated on the basis of known marker genes and those sharing markers (e.g. olfactory sensory neurons) were merged.

The mouse WOM Drop-seq dataset contained 29585 cells that passed the above filtering. Each of the 16 clusters identified contained cells from all 8 replicates in roughly equal proportions. Of the 17666 mature OSNs and the 4674 immature OSNs, none of the cells express Ace2. In contrast, in the olfactory epithelial cells, Ace2 expression was observed in the Bowman’s gland, olfactory HBCs, dorsal sustentacular cells.

### Immunohistochemistry and in situ hybridization of mouse and human tissue

#### Mouse olfactory epithelium tissue processing

Mice were sacrificed with a lethal dose of xylazine and nasal epithelium with attached olfactory bulbs were dissected and fixed in 4% paraformaldehyde (Electron Microscope Sciences, 19202) in phosphate-buffered saline (PBS) for overnight at 4°C or for 2 hours at room temperature. Tissues were washed in PBS for 3 times (5 min each) and incubated in 0.45M EDTA in PBS overnight at 4°C. The following day, tissues were rinsed by PBS and incubated in 30 % Sucrose in PBS for at least 30 min, transferred to Tissue Freezing Medium (VWR, 15146-025) for at least 45 min and frozen on crushed dry ice and stored at −80°C until sectioning. Tissue sections (20 μm thick for the olfactory bulb and 12 μm thick for nasal epithelium) were collected on Superfrost Plus glass slides (VWR, 48311703) and stored at −80°C until immunostaining.

For methimazole treated samples, Adult C57BL/6J mice (6-12 weeks old, JAX stock No. 000664) were given intraperitoneal injections with Methimazole (Sigma M8506) at 50 μg/g body weight and sacrificed at 24, 48, and 96-hour timepoints.

#### Immunostaining for mouse tissue

Sections were permeabilized with 0.1% Triton X-100 in PBS for 20 min then rinsed 3 times in PBS. Sections were then incubated for 45-60 min in blocking solution that consisted of PBS containing 3% Bovine Serum Albumin (Jackson Immunoresearch, 001-000-162) and 3% Donkey Serum (Jackson ImmunoResearch, 017-000-121) at room temperature, followed by overnight incubation at 4°C with primary antibodies diluted in the same blocking solution. Primary antibodies used are as follows. Goat anti-ACE2 (Thermo Fisher, PA5-47488, 1:40), mouse anti-TUBB4 (Sigma, T7941, 1:4000), rabbit anti-KRT5 (abcam, ab52635, 1:200), goat anti-NQO1 (abcam, ab2346, 1:200), mouse anti-acetylated Tubulin (abcam, ab24610, 1:500), rabbit anti-CNGA2 (abcam, ab79261, 1:100), rat anti-CD140b/PDGFRB (Thermo Fisher, 14-1402-82, 1:100).

On the following day, sections were rinsed once and washed three times for 5-10 min in PBS, then incubated for 45 min with secondary antibodies diluted in blocking solution at 1:300 ratios and/or Alexa 555-conjugated Phalloidin (1:400). Secondary antibodies used were as follows: Donkey anti-Goat IgG Alexa 488 (Jackson ImmunoResearch, 705-546-147), donkey anti-Goat IgG Alexa 555, (Invitrogen, A21432), donkey anti-Rabbit IgG Alexa 555 (Invitrogen, A31572), donkey anti-Rabbit IgG Alexa 647 (Jackson ImmunoResearch, 711-605-152), donkey anti-Mouse IgG Alexa 555 (Invitrogen, A31570), donkey anti-Mouse IgG Alexa 647 (Invitrogen, A31571), and donkey anti-Rat IgG Alexa 488 (Invitrogen, A21208).

After secondary antibody incubation, sections were washed twice for 5-10 min in PBS, incubated with 300 nM DAPI in PBS for 10 min and then rinsed with PBS. Slides were mounted with glass coverslips using Vectashield Mounting Medium (Vector Laboratories, H-1000) or ProLong Diamond Antifade Mountant (Invitrogen, P36961).

For co-staining of ACE2 and NQO1, slides were first stained with ACE2 primary antibody and donkey anti-Goat IgG Alexa 488 secondary. After 3 washes of secondary antibody, tissues were incubated with unconjugated donkey anti-Goat IgG Fab fragments (Jackson ImmunoResearch, 705-007-003) at 30 μg/mL diluted in blocking solution for 1 hour at room temperature. Tissues were washed twice with PBS, once in blocking solution, and incubated in blocking solution for 30-40 min at room temperature, followed by a second round of staining with the NQO1 primary antibody and donkey anti-Goat IgG Alexa 555 secondary antibody.

Confocal images were acquired using a Leica SPE microscope (Harvard Medical School Neurobiology Imaging Facility) with 405 nm, 488 nm, 561 nm, and 635 nm laser lines. Multi-slice z-stack images were acquired, and their maximal intensity projections are shown. For Figure 3E, tiled images were acquired and stitched by the Leica LAS X software. Images were processed using Fiji ImageJ software (*60*), and noisy images were median-smoothed using the *Remove Outliers* function built into Fiji.

#### Fluorescent in situ hybridization for mouse tissue

Sult1c1 RNA was detected by fluorescent RNAscope assay (Advanced Cell Diagnostics, kit 320851) using probe 539921-C2, following the manufacturer’s protocol (RNAscope Fluorescent Multiplex Kit User Manual, 320293-UM Date 03142017) for paraformaldehyde-fixed tissue. Prior to initiating the hybridization protocol, the tissue was pre-treated with two successive incubations (first 30 min, then 15 min long) in RNAscope Protease III (Advanced Cell Diagnostics, 322337) at 40°C, then washed in distilled water. At the end of protocol, the tissue was washed in PBS and subjected to the 2-day immunostaining protocol described above.

#### Immunostaining of human nasal tissue

Human olfactory mucosa biopsies were obtained via IRB-approved protocol at Duke University School of Medicine, from nasal septum or superior turbinate during endoscopic sinus surgery. Tissue was fixed with 4% paraformaldehyde and cryosectioned at 10 μm and sections were processed for immunostaining, as previously described (*37*).

Sections from a female nasal septum biopsy were stained for ACE2 (Figure 2F) using the same Goat anti-ACE2 (Thermo Fisher, PA5-47488, 1:40) and the protocol described above for mouse tissue. The human sections were co-stained with Rabbit anti-keratin 5 (Abcam, ab24647; AB_448212, 1:1000) and were detected with AlexaFluor 488 Donkey anti-goat (Jackson ImmunoResearch, 705-545-147) and AlexaFluor 594 Donkey anti-rabbit (Jackson ImmunoResearch, 711-585-152) secondary antibodies (1:300).

As further validation of ACE2 expression and to confirm the lack of ACE2 expression in human olfactory sensory neurons (Figure S1E), sections were stained with a rabbit anti-ACE2 (Abcam, ab15348; RRID:AB_301861, used at 1:100) antibody immunogenized against human ACE2 and a mouse Tuj1 antibody against neuron-specific tubulin (BioLegend, 801201; RRID:AB_2313773). Anti-ACE2 was raised against a C-terminal synthetic peptide for human ACE2 and was validated by the manufacturer to not cross-react with ACE1 for immunohistochemical labeling of ACE2 in fruit bat nasal tissue as well as in human lower airway. Recombinant human ACE2 abolished labeling with this antibody in a previous study in human tissue, further demonstrating its specificity (*61*). The Tuj1 antibody was validated, as previously described (*37*). Biotinylated secondary antibodies (Vector Labs), avidin-biotinylated horseradish peroxidase kit (Vector) followed by fluorescein tyramide signal amplification (Perkin Elmer) were applied per manufacturer’s instructions. For dual staining, Tuj1 was visualized using AlexaFluor 594 Goat anti-Mouse (Jackson ImmunoResearch, 115-585-146; RRID: AB_2338881).

Human sections were counterstained with 4’,6-diamidino-2-phenylindole (DAPI) and coverslips were mounted using Prolong Gold (Invitrogen) for imaging, using a Leica DMi8 microscope system. Images were processed using Fiji ImageJ software (NIH). Scale bars were applied directly from the Leica acquisition software metadata in ImageJ Tools. Unsharp Mask was applied in ImageJ, and brightness/contrast was adjusted globally.

### WOM and HBC lineage tracing mouse 10x scSeq experiments

#### Mice

2 month-old and 18 month-old wild type C57BL/6J mice were obtained from the National Institute on Aging Aged Rodent Colony and used for the WOM experiments; each experimental condition consisted of one male and one female mouse to aid doublet detection. Mice containing the transgenic Krt5-CreER(T2) driver (*62*) and Rosa26-YFP reporter allele (*63*) were used for the HBC lineage tracing dataset. All mice were assumed to be of normal immune status. Animals were maintained and treated according to federal guidelines under IACUC oversight at the University of California, Berkeley.

#### Single-Cell RNA Sequencing

The olfactory epithelium was surgically removed, and the dorsal, sensory portion was dissected and dissociated, as previously described (*32*). For WOM experiments, dissociated cells were subjected to fluorescence-activated cell sorting (FACS) using propidium iodide to identify and select against dead or dying cells; 100,000 cells/sample were collected in 10% FBS. For the HBC lineage tracing experiments Krt5-CreER; Rosa26YFP/YFP mice were injected once with tamoxifen (0.25 mg tamoxifen/g body weight) at P21-23 days of age and sacrificed at 24 hours, 48 hours, 96 hours, 7 days and 14 days post-injury, as previously described (*32, 64*). For each experimental time point, YFP+ cells were isolated by FACS based on YFP expression and negative for propidium iodide, a vital dye.

Cells isolated by FACS were subjected to single-cell RNA-seq. Three replicates (defined here as a FACS collection run) per age were analyzed for the WOM experiment; at least two biological replicates were collected for each experimental condition for the HBC lineage tracing experiment. Single cell cDNA libraries from the isolated cells were prepared using the Chromium Single Cell 3’ System according to the manufacturer’s instructions. The WOM preparation employed v3 chemistry with the following modification: the cell suspension was directly added to the reverse transcription master mix, along with the appropriate volume of water to achieve the approximate cell capture target. The HBC lineage tracing experiments were performed using v2 chemistry. The 0.04% weight/volume BSA washing step was omitted to minimize cell loss. Completed libraries were sequenced on Illumina HiSeq4000 to produce paired-end 100nt reads.

Sequence data were processed with the 10x Genomics Cell Ranger pipeline (2.0.0 for v2 chemistry), resulting in the initial starting number before filtering of 60,408 WOM cells and 25,469 HBC lineage traced cells. The scone R/Bioconductor package (*65*) was used to filter out lowly-expressed genes (fewer than 2 UMI’s in fewer than 5 cells) and low-quality libraries (using the metric_sample_filter function with arguments hard_nreads = 2000, zcut = 4).

#### Preliminary Filtering

Cells with co-expression of male (Ddx3y, Eif2s3y, Kdm5d, and Uty) and female marker genes (Xist) were removed as potential doublets from the WOM dataset. For both datasets, doublet cell detection was performed per sample using DoubletFinder (*66*) and Scrublet (*67*). Genes with at least 3 UMIs in at least 5 cells were used for downstream clustering and cell type identification. For the HBC lineage tracing dataset, the Bioconductor package scone was used to pick the top normalization (“none,fq,ruv_k=1,no_bio,batch”), corresponding to full quantile normalization, batch correction and removing one factor of unwanted variation using RUV (*68*). A range of cluster labels were created by clustering using the partitioning around medoids (PAM) algorithm and hierarchical clustering in the clusterExperiment Bioconductor package (*69*), with parameters k0s=(10,13,16,19,22,25) and alpha=(NA,0.1,0.2,0.3). Clusters that did not show differential expression were merged (using the function mergeClusters with arguments mergeMethod = ‘adjP’, cutoff = 0.01, and DEMethod = ‘limma’ for the lineage-traced dataset). Initial clustering identified one Macrophage (Msr1+) cluster consisting of 252 cells; upon its removal and restarting from the normalization step a subsequent set of 15 clusters was obtained. These clusters were used to filter out 1515 cells for which no stable clustering could be found (i.e., ‘unassigned’ cells), and four clusters respectively consisting of 31, 29 and 23 and 305 cells. Doublets were identified using DoubletFinder and 271 putative doublets were removed. Inspection of the data in a three-dimensional UMAP embedding identified two groups of cells whose experimentally sampled timepoint did not match their position along the HBC differentiation trajectory, and these additional 219 cells were also removed from subsequent analyses.

#### Analysis of CoV-related genes in WOM and HBC lineage 10x datasets

Analysis of WOM scSeq data were performed in python using the open-source Scanpy software starting from the raw UMI count matrix of the 40179 cells passing the initial filtering and QC criteria described above. UMIs were total-count normalized and scaled by 10,000 (TPT, tag per ten-thousands) and then log-normalized. For each gene, the residuals from linear regression models using the total number of UMIs per cell as predictors were then scaled via z-scoring. PCA was then performed on a set of highly-variable genes (excluding OR genes) calculated using the “highly_variable_genes” function with parameters: min_mean=0.01, max_mean=10, min_disp=0.5. A batch corrected neighborhood graph was constructed by the “bbknn” function with 42 PCs with the parameters: local_connectivity=1.5, and embedding two-dimensions using the UMAP function with default parameters (min_dist = 0.5). Cells were clustered using the neighborhood graph via the Leiden algorithm (resolution = 1.2). Identified clusters were manually merged and annotated based on known marker gene expression. We removed 281 cells containing mixtures of marker genes with no clear gene expression signature. The identified cell types and the number of each of the remaining 39898 cells detected were as follows. 28,769 mOSN: mature OSN; 2,607 iOSN: immature OSN; 859 INP: Immediate Neural Precursor; 623 GBC: Globose Basal Cell; HBC: Horizontal Basal Cell (1,083 Olfactory and 626 Respiratory); 480 SUS: sustentacular cell; 331 BG: Bowman’s gland; MV: Microvillar cell (563 Brush-like and 1,530 Ionocyte-like); 92 OEC: Olfactory Ensheathing Cell; 76 Resp. Secretory cells; 227 Resp. unspecified cells; 172 atypical OSN; 1,757 various immune cells, 103 RBC: Red Blood Cell. TPT gene expression levels were visualized in two-dimensional UMAP plots.

The filtered HBC lineage dataset containing 21722 cells was analyzing in python and processed for visualization using pipelines in SPRING and Scanpy (*70, 71*). In brief, total counts were normalized to the median total counts for each cell and highly variable genes were selected using the SPRING gene filtering function (“filter_genes”) using parameters (90, 3, 3). The dimensionality of the data was reduced to 20 using principal components analysis (PCA) and visualized in two-dimensions using the UMAP method with parameters (n_neighbors=20, min_dist=0.5). Clustering was performed using the Leiden algorithm (resolution=1.45) and clusters were merged manually using known marker genes. The identified cell types and number of each type were: 929 mOSN: mature OSN; 2073 iOSN: immature OSN; 786 INP: Immediate Neural Precursor; 755 GBC: Globose Basal Cell; HBC: Horizontal Basal Cell (7782 Olfactory, 5418 Regenerating, and 964 Respiratory); 2666 SUS: sustentacular cell; and 176 Ionocyte-like Microvillar (MV) cell.

Expression of candidate CoV-2-related genes was defined if at least one transcript (UMI) was detected in that cell, and the percent of cells expressing candidate genes was calculated for each cell type. In the WOM dataset Ace2 was only detected in 2 out of 28,769 mature OSNs (0.007 %), and in the HBC lineage dataset, Ace2 was not detected in any OSNs. Furthermore, Ace2 was not detected in immature sensory neurons (GBCs, INPs, or iOSNs) in either dataset.

### Mouse HBC lineage Smart-Seq2 dataset

Single-cell RNA-seq data from HBC-derived cells from Fletcher et al. and Gadye et al (*32, 64*), labeled via *Krt5-CreER* driver mice, were downloaded from GEO at accession GSE99251 using the file “GSE95601_oeHBCdiff_Cufflinks_eSet_counts_table.txt.gz”. Processing was performed as described above, including total counts normalization and filtering for highly variable genes using the SPRING gene filtering function “filter_genes” with parameters (75, 20, 10). The resulting data were visualized in SPRING and a subset of cells were removed for quality control, including a cluster of cells with low total counts and another with predominantly reads from ERCC spike-in controls. Putative doublets were also identified using Scrublet and removed (6% of cells) (*67*). The resulting data were visualized in SPRING and partitioned using Louvain clustering on the SPRING k-nearest-neighbor graph using the top 40 principal components. Cell type annotation was performed manually using the same set of markers genes listed above. Three clusters were removed for quality control, including one with low total counts and one with predominantly reads from ERCC spike-in controls (likely background), and one with high mitochondrial counts (likely stressed cells). For visualization, and clustering the remaining cells were projected to 15 dimensions using PCA and visualized with UMAP with parameters (n_neighbors=15, min_dist=0.4, alpha=0.5, maxiter=500). Clustering was performed using the Leiden algorithm (resolution=0.4) and cell types were manually annotated using known marker genes.

The filtered dataset of mouse HBC-derived cells contained 1450 cells. The percent of cells expressing each marker gene was calculated as described above. Of the 51 OSNs identified, none of them expressed Ace2, and only 1 out of 194 INPs and iOSNs expressed Ace2. In contrast, Ace2 and Tmprss2 were both detected in HBCs and SUS cells.

### Juvenile and adult mouse whole olfactory bulb scRNAseq dataset

#### Juvenile mouse data

Single-cell RNAseq data from whole mouse olfactory bulb (*42*) were downloaded from mousebrain.org/loomfiles_level_L1.html in loom format (l1 olfactory.loom) and converted to a Seurat object. Samples were obtained from juvenile mice (age postnatal day 26-29). This dataset comprises 20514 cells passing cell quality filters, excluding 122 cells identified as potential doublets.

#### Tissue dissection and single-cell dissociation

A new dataset of whole olfactory bulb scSeq was generated from adult male mice (8–12 weeks-old). All mouse husbandry and experiments were performed following institutional and federal guidelines and approved by Harvard Medical School’s Institutional Animal Care and Use Committee (IACUC). Briefly, dissected olfactory bulbs (including the accessory olfactory bulb and fractions of the anterior olfactory nucleus) were dissociated in 750 μl of dissociation media (DM: HBSS containing 10mM HEPES, 1 mM MgCl2, 33 mM D-glucose) with 28 U/mL Papain and 386 U/mL DNase I (Worthington). Minced tissue pieces were transferred to a 5 mL round-bottom tube (BD). DM was added to a final volume of 3.3 mL and the tissue was mechanically triturated 5 times with a P1000 pipette tip. After 1-hour incubation with rocking at 37°C, the suspension was triturated with a 10 mL pipette 10 times and 2.3 mL was passed through 40 μm cell strainer (BD). The suspension was then mechanically triturated with a P1000 pipette tip 10 times and 800 μL were filtered on the same strainer. The cell suspension was further triturated with a P200 pipette tip 10 times and filtered. 1 mL of Quench buffer (22 mL of DM, 2.5 mL of protease inhibitor prepared by resuspending 1 vial of protease inhibitor with 32 mL of DM, and 2000U of DNase I) was added to the suspension and centrifuged at 300g for 5 min. Cells were resuspended with 3 mL of Quench buffer and overlaid gently on top of 5 mL of protease inhibitor, then spun down at 70g for 10min. The pellet was resuspended using DM supplemented with 0.04 % BSA and spun down at 300g for 5 min. Cells were suspended in 400 μL of DM with 0.04 % BSA.

#### Olfactory bulb Drop-seq experiments

Drop-seq experiments were performed as previously described (*59*). Microfluidics devices were obtained from FlowJEM and barcode beads were obtained from chemgenes. Two 15 min Drop-seq runs were collected from a single dissociation preparation obtained from 2 mice. Two such dissociations were performed, giving 4 total replicates.

#### Sequencing of Drop-seq samples

4 replicates of Drop-seq samples were pooled and sequenced across 3 runs on an Illumina NextSeq 500 platform. Paired end reads from the fastq files were trimmed, aligned, and tagged via the Drop-seq tools (1-2.0) pipeline, using STAR (2.4.2a) with genomic indices from Ensembl Release 82. The digital gene expression matrix was generated for 8,000 cells per replicate.

#### Preprocessing of Drop-seq samples

Cells with low numbers of genes (500), low numbers of UMIs (700) or high numbers of UMIs (>10000) were removed (6 % of cells). Potential doublets were identified via Scrublet and removed (3.5 % of cells). Overall, this new dataset comprised 27004 cells.

#### Integration of whole olfactory bulb scRNAseq datasets

Raw UMI counts from juvenile and adult whole olfactory bulb samples were integrated in Seurat (*72*). Integrating the datasets ensured that clusters with rare cell types could be identified and that corresponding cell types could be accurately matched. As described below (see Figure S5), although some cell types were observed with different frequencies, the integration procedure yielded stable clusters with cells from both datasets. Briefly, raw counts were log-normalised separately and the 10000 most variable genes identified by variance stabilizing transformation for each dataset. The 4529 variable genes present in both datasets and the first 30 principal components (PCs) were used as features for identifying the integration anchors. The integrated expression matrix was scaled and dimensionality reduced using PCA. Based on their percentage of explained variance, the first 28 PCs were chosen for UMAP visualisation and clustering.

Graph-based clustering was performed using the Louvain algorithm following the standard Seurat workflow. Cluster stability was analysed with Clustree on a range of resolution values (0.4 to 1.4), with 0.6 yielding the most stable set of clusters (*73*). Overall, 26 clusters were identified, the smallest of which contained only 43 cells with gene expression patterns consistent with blood cells, which were excluded from further visualisation plots. Clustering the two datasets separately yielded similar results. Moreover, the distribution of cells from each dataset across clusters was homogenous (Figure S5) and the clusters corresponded previous cell class and subtype annotations (*42*). As previously reported, a small cluster of excitatory neurons (cluster 13) contained neurons from the anterior olfactory nucleus. UMAP visualisations of expression level for cell class and cell type markers, and for genes coding for coronavirus entry proteins, depict log-normalized UMI counts. The heatmap in Figure 4B shows the mean expression level for each cell class, normalised to the maximum mean value. The percentage of cells per cell class expressing Ace2 was defined as the percentage of cells with at least one UMI. In cells from both datasets, Ace2 was enriched in pericytes but was not detected in neurons.

### Smart-Seq2 sequencing of manually sorted olfactory bulb dopaminergic neurons

#### Tissue dissociation and manual cell sorting

Acute olfactory bulb 300 μm slices were obtained from Dat-Cre/Flox-tdTomato (B6.SJL-Slc6a3^tm1.1(cre)Bkmn/J^, Jax stock 006660 / B6.Cg–Gt(ROSA)26Sor^tm9(CAG-tdTomato)Hze^, Jax stock 007909) P28 mice as previously described (*74*). As part of a wider study, at P27 these mice had undergone brief 24 h unilateral naris occlusion via a plastic plug insert (N = 5 mice) or were subjected to a sham control manipulation (N = 5 mice); all observed effects here were independent of these treatment groups. Single cell suspensions were generated using the Neural Tissue Dissociation Kit – Postnatal Neurons (Miltenyi Biotec. Cat no. 130-094-802), following manufacturer’s instructions for manual dissociation, using 3 fired-polished Pasteur pipettes of progressively smaller diameter. After enzymatic and mechanical dissociations, cells were filtered through a 30 μm cell strainer, centrifuged for 10 minutes at 4° C, resuspended in 500 μl of ACSF (in mM: 140 NaCl, 1.25 KCl, 1.25 NaH_2_PO_4_, 10 HEPES, 25 Glucose, 3 MgCl_2_, 1 CaCl_2_) with channel blockers (0.1 μM TTX, 20 μM CNQX, 50 μM D-APV) and kept on ice to minimise excitotoxicity and cell death.

For manual sorting of fluorescently labelled dopaminergic neurons we adapted a previously described protocol (*75*). 50 μl of single cell suspension was dispersed on 3.5mm petri dishes (with a Sylgard-covered base) containing 2 ml of ACSF + channel blockers. Dishes were left undisturbed for 15 minutes to allow the cells to sink and settle. Throughout, dishes were kept on a metal plate on top of ice. tdTomato-positive cells were identified by their red fluorescence under a stereoscope. Using a pulled glass capillary pipette attached to a mouthpiece, individual cells were aspirated and transferred to a clean, empty dish containing 2 ml ACSF + channel blockers. The same cell was then transferred to a third clean plate, changing pipettes for every plate change. Finally, each individual cell was transferred to a 0.2 ml PCR tube containing 2 μl of lysis buffer (RLT Plus - Qiagen). The tube was immediately placed on a metal plate sitting on top of dry ice for flash-freezing. Collected cells were stored at −80C until further processing. Positive (more than 10 cells) and negative (sample collection procedure without picking a cell) controls were collected for each sorting session. In total, we collected samples from 10 mice, averaging 50 tdTomato-positive cells collected per session. Overall, less than 2.5 hours elapsed between mouse sacrifice and collection of the last cell in any session.

#### Preparation and amplification of full-length cDNA and sequencing libraries

Samples were processing using a modified version of the Smart-Seq2 protocol(*76*). Briefly, 1 μl of a 1:2,000,000 dilution of ERCC spike-ins (Invitrogen. Cat. no. 4456740) was added to each sample and mRNA was captured using modified oligo-dT biotinylated beads (Dynabeads, Invitrogen). PCR amplification was performed for 22 cycles. Amplified cDNA was cleaned with a 0.8:1 ratio of Ampure-XP beads (Beckman Coulter). cDNAs were quantified on Qubit using HS DNA reagents (Invitrogen) and selected samples were run on a Bioanalyzer HS DNA chip (Agilent) to evaluate size distribution.

For generating the sequencing libraries, individual cDNA samples were normalised to 0.2ng/μl and 1μl was used for one-quarter standard-sized Nextera XT (Illumina) tagmentation reactions, with 12 amplification cycles. Sample indexing was performed using index sets A and D (Illumina). At this point, individual samples were pooled according to their index set. Pooled libraries were cleaned using a 0.6:1 ratio of Ampure beads and quantified on Qubit using HS DNA reagents and with the KAPA Library Quantification Kits for Illumina (Roche). Samples were sequenced on two separate rapid-runs on HiSeq2500 (Illumina), generating 100bp paired-end reads. An additional 5 samples were sequenced on MiSeq (Illumina).

#### Full-length cDNA sequencing data processing and analysis

Paired-end read fastq files were demultiplexed, quality controlled using FastQC (https://www.bioinformatics.babraham.ac.uk/projects/fastqc/) and trimmed using Trim Galore (https://www.bioinformatics.babraham.ac.uk/projects/trim_galore/). Reads were pseudoaligned and quantified using kallisto (*77*) against a reference transcriptome from Ensembl Release 89 (Gencode Release M17 GRCm38.p6) with sequences corresponding to the ERCC spike-ins and the Cre recombinase and tdT genes added to the index. Transcripts were collapsed into genes using the sumAcrossFeatures function in scater.

Cell level quality control and cell filtering was performed in scater (*78*). Cells with <1000 genes, <100,000 reads, >75% reads mapping to ERCC spike-ins, >10% reads mapping to mitochondrial genes or low library complexity were discarded (14% samples). The population of olfactory bulb cells labelled in DAT-tdTomato mice is known to include a minor non-dopaminergic calretinin-positive subgroup (*79*), so calretinin-expressing cells were excluded from all analyses. The scTransform function in Seurat was used to remove technical batch effects.

### Expression of CoV-relevent genes in scSeq datasets from various brain regions and sensory systems

An analysis of single-cell gene expression data from 10 studies was performed to investigate the expression of genes coding for coronavirus entry proteins in neurons from a range of brain regions and sensory systems. Processed gene expression data tables were obtained from scSeq studies that evaluated gene expression in retina (GSE81905) (*80*) inner ear sensory epithelium (GSE115934) (*81, 82*) and spiral ganglion (GSE114997) (*83*), ventral midbrain (GSE76381) (*84*), hippocampus (GSE100449) (*85*), cortex (GSE107632) (*86*), hypothalamus (GSE74672) (*87*), visceral motor neurons (GSE78845) (*88*), dorsal root ganglia (GSE59739) (*89*) and spinal cord dorsal horn (GSE103840) (*90*). Smart-Seq2 sequencing data from Vsx2-GFP positive cells was used from the retina dataset. A subset of the expression matrix that corresponds to day 0 (i.e. control, undisturbed neurons) was used from the layer VI somatosensory cortex dataset. A subset of the data containing neurons from untreated (control) mice was used from the hypothalamic neuron dataset. From the ventral midbrain dopaminergic neuron dataset, a subset comprising DAT-Cre/tdTomato positive neurons from P28 mice was used. A subset comprising Type I neurons from wild type mice was used from the spiral ganglion dataset. The “unclassified” neurons were excluded from the visceral motor neuron dataset. A subset containing neurons that were collected at room temperature was used from the dorsal root ganglia dataset. Expression data from dorsal horn neurons obtained from C57/BL6 wild type mice, vGat-cre-tdTomato and vGlut2-eGFP mouse lines was used from the spinal cord dataset. Inspection of all datasets for batch effects was performed using the scater package (version 1.10.1) (*78*). Publicly available raw count expression matrices were used for the retina, hippocampus, hypothalamus, midbrain, visceral motor neurons and spinal cord datasets, whereas the normalized expression data was used from the inner ear hair cell datasets. For datasets containing raw counts, normalization was performed for each dataset separately by computing pool-based size factors that are subsequently deconvolved to obtain cell-based size factors using the scran package (version 1.10.2) (*91*). Violin plots were generated in scater.

## References and Notes

## Acknowledgments

We thank members of the Datta lab, James Schwob, Bernardo Sabatini, Andreas Schaefer, Kevin Franks, Michael Greenberg and Vanessa Ruta for helpful comments on the manuscript. We thank James Lipscombe and Andres Crespo for technical support.

## Author contributions

DHB, TT, RC, HC, RF, YC, ML, ICM, RAH and BJG designed and performed experiments. DHB, TT, CW, ML, KvdB, BG, RC, HC, DD, KS, and HRdB performed analysis. DR, SD, EP, JSM, BJG, MSG JN, SRD, designed and supervised experiments. DHB, TT, CW, ML, HM, DWL, BJG, MSG, JN and SRD wrote the manuscript

## Data and materials availability

Reanalyzed datasets are obtained from the URLs listed in supplementary materials. All data is currently being deposited and will be made publicly accessible from the NCBI GEO at accession GSE148360.

## Supplementary Materials

### URLs for re-analyzed datasets

https://www.ncbi.nlm.nih.gov/pmc/articles/PMC4680959/bin/srep18178-s2.xls

https://doi.org/10.1371/journal.pone.0113170.s014

https://www.ncbi.nlm.nih.gov/geo/query/acc.cgi?acc=GSE52464

https://advances.sciencemag.org/highwire/filestream/217162/field_highwire_adjunct_files/0/aax0396_Data_file_S1.xlsx

https://www.ncbi.nlm.nih.gov/geo/query/acc.cgi?acc=GSE139522

https://www.ncbi.nlm.nih.gov/geo/query/acc.cgi?acc=GSE99251

https://www.ncbi.nlm.nih.gov/geo/query/acc.cgi?acc=GSE120199

https://www.genomique.eu/cellbrowser/HCA/

**Fig. S1.**
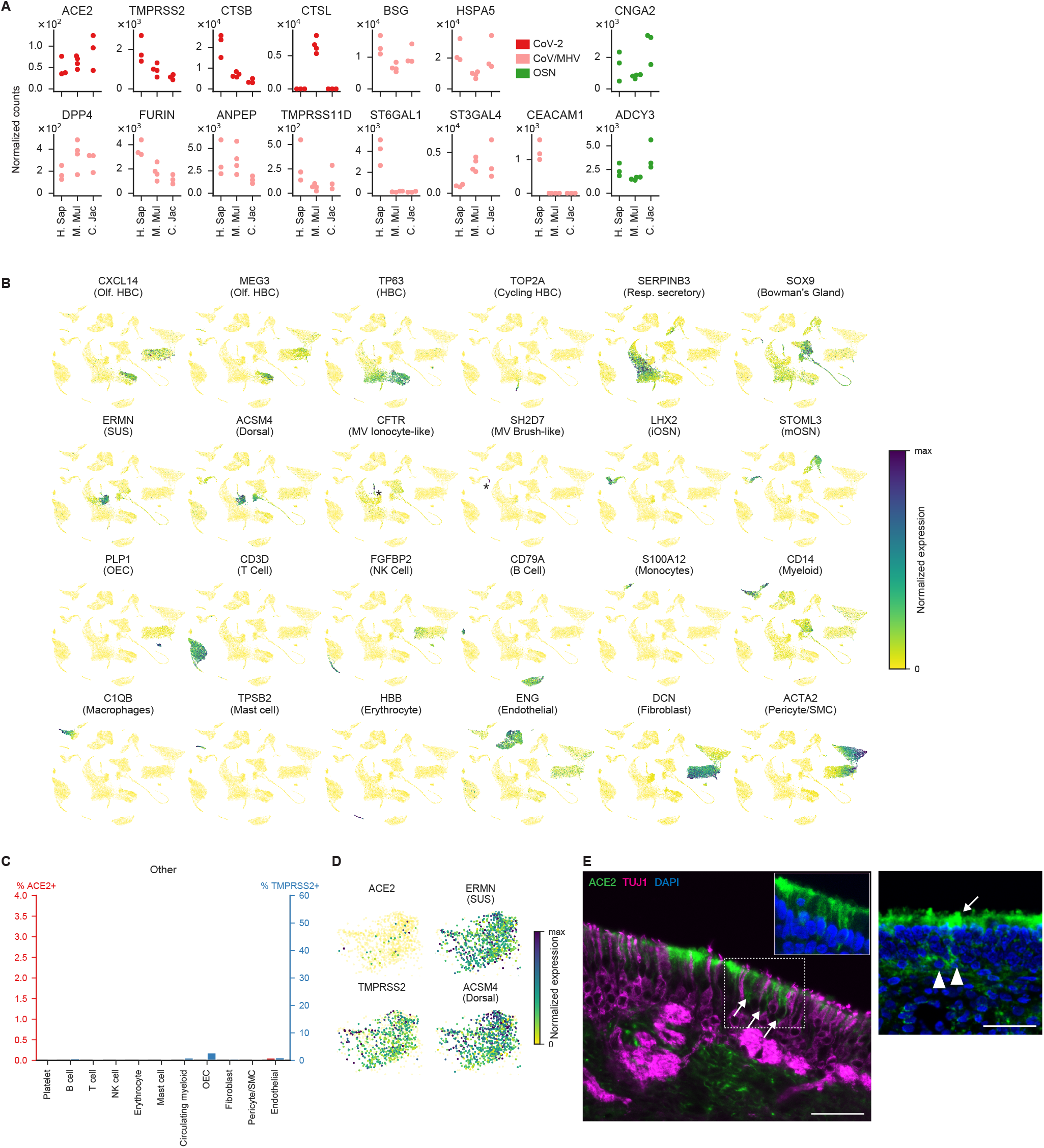
Related to Fig. 1. Bulk RNA-seq datasets for primates and analysis of human scSeq data. **(A)** Expression of genes required for the entry of coronavirus (CoV) and OSN markers in primate olfactory mucosa in the data from Saraiva et al 2019 (*36*). Human (H. Sap), Macaque (M. Mul) and Marmoset (C. Jal) data are shown. Each circle represents a biological replicate and each color indicates the category of the gene shown on the right. Raw counts were normalized to account for differences in sequencing depth between samples. **(B)** UMAP representation of the mouse WOM Drop-seq dataset shown in Figure 3, with normalized expression for all the cell types shown in Fig. 3B indicated (HBC = horizontal basal cells, Resp = respiratory, SUS = sustentacular cells, MV = microvillar cells, iOSN = immature olfactory sensory neurons, mOSNs = mature olfactory sensory neurons, OEC = olfactory ensheathing cells, SMC = smooth muscle cells). CFTR-expressing MV Ionocyte-like cells and SH2D7-expressing MV Brush-like cells are indicated by asterisks. **(C)** Percent of cells expressing ACE2 and TMPRSS2 in the cell types not shown in Fig. 2B and plotted on the same scale as Fig. 3D. ACE2 was rarely detected in non-epithelial cell types. **(D)** UMAP representations of all sustentacular (SUS) cells with the normalized expression of CoV-2 related genes ACE2 and TMPRSS2, as well as sustentacular (ERMN) cell markers. The majority of SUS cells captured in (*37*), including the ACE2-positive ones, expressed dorsal cell (ACSM4) markers. **(E)** ACE2 immunostaining of human olfactory mucosal biopsy samples, using a different antibody than shown in **Figure 2** for validation purposes. ACE2 protein (green) is detected in sustentacular cells (white arrows) in both 86-year old male sample (left) and 39-year old female sample (right). Inset in the left image is an enlarged view of the dashed box showing ACE2 and nuclei, instead of TUJ1, for clear visualization of ACE2 signal. ACE2 does not appear to colocalize with OSN marker TUJ1 (magenta). ACE2 is also detected in basal cells (white arrowheads in the middle image). Nuclei were stained with DAPI. Bar = 50 μm.

**Fig. S2.**
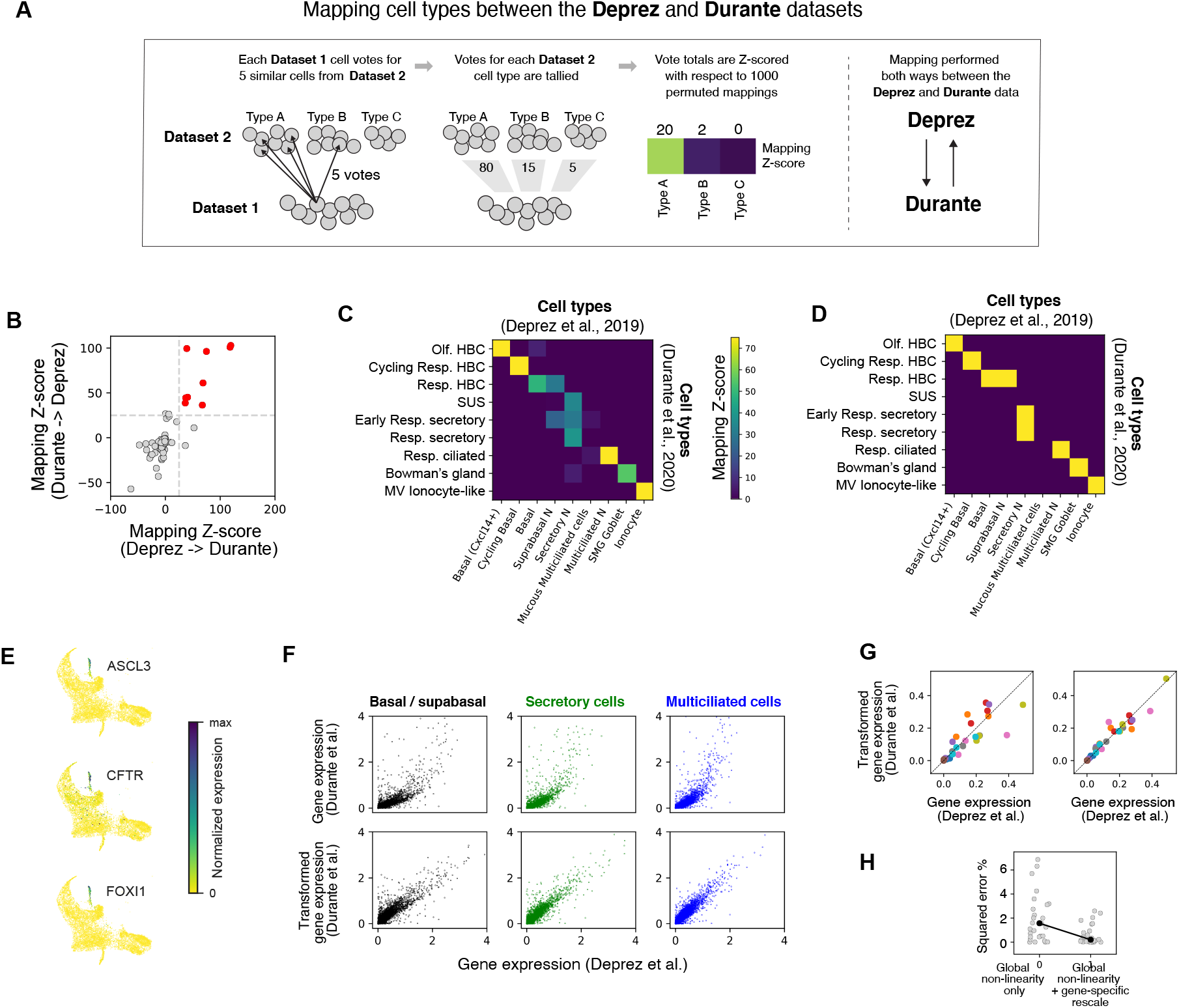
Related to Fig. 2. Comparing cell types and expression levels across respiratory and olfactory epithelial datasets. **(A)** Schematic of the mapping strategy used to identify similar cell types across datasets, applied to a toy example. Each cell type from “Dataset 1” dataset is mapped to cell types from the “Dataset 2”. From left to right: Each Dataset 1 cell voted on its 5 most similar cells in Dataset 2; the total number of votes cast for each Dataset 2 cell type was quantified; and vote totals were Z-scored against 1000 shuffles where cell type labels were permutated. **(B)** Mapping was performed bi-directionally between the Deprez and Durante datasets, and the mapping Z-scores in each direction are compared. The set of cell type correspondences with high Z-scores (>25) in both directions are colored red. **(C)** Mapping Z-scores from one of the two mappings (Deprez -> Durante). **(D)** The set of cell type correspondences with high bi-directional mappings (both Z-scores > 25; equivalent to the red dots in **B**). **(E)** Gene expression in olfactory microvillar cells from the Durante dataset. These cells express classical microvillar genes (ASCL3) as well as marker genes of pulmonary ionocytes (CFTR, FOXI1). **(F)** Top: average expression for each gene in three cell types compared between the Durante and Deprez datasets in the units of the original papers. Bottom: average expression after a global non-linear transformation of the Durante data. Basal/suprabasal cells = “respiratory HBCs” from Durante et al., and “basal” and “suprabasal” cells from Deprez et al. Secretory cells = “early respiratory secretory cells” and “respiratory secretory cells” from Durante et al., and “secretory” cells from Deprez et al. Mutliciliated cells = “respiratory ciliated cells” from Durante et al., and “multiciliated” cells from Deprez et al. **(G)** Average expression of genes shown in Fig 3G before (left) and after (right) gene-specific rescaling. Each color is one gene. For every color, three dots are shown, corresponding to expression across the three cell types in **F**. The diagonal line represents unity. Dots close to the diagonal line indicate a close match in transformed gene expression the Durante and Deprez data for the cell types shown in **F**. **(H)** Comparison of percent squared error before versus after gene-specific rescaling.

**Fig. S3.**
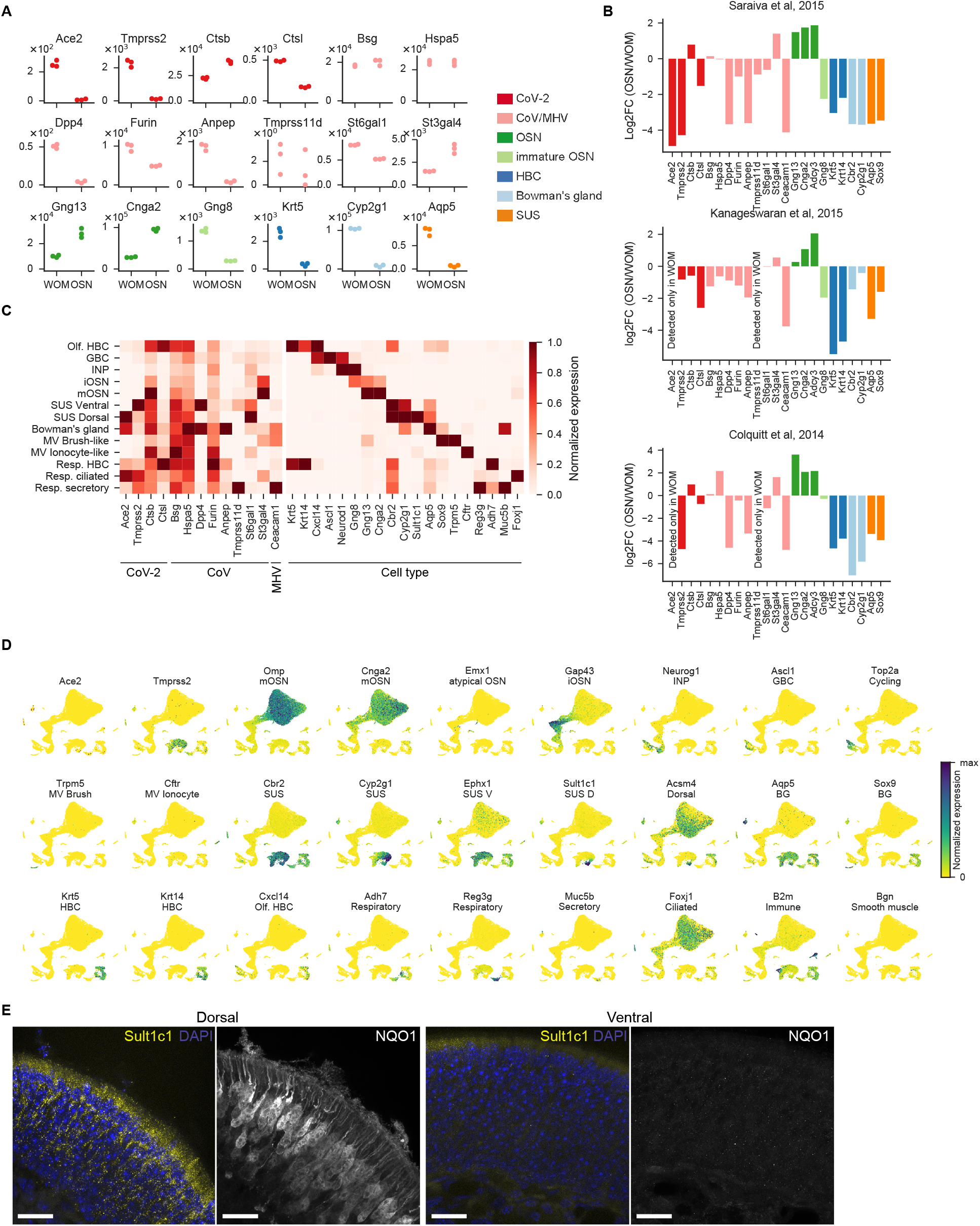
Related to Fig. 2. Analysis of mouse bulk RNA-Seq, a WOM scSeq (Drop-seq) dataset and validation of dorsal sustentacular cell marker Sult1c1. **(A)** Expression of coronavirus (CoV)-related genes and cell type markers in mouse olfactory mucosa from Saraiva et al. 2015 (*39*). Normalized counts for each gene in the whole olfactory mucosa (WOM) and olfactory sensory neurons (OSNs) are shown. Each circle represents a biological replicate and each color indicates the category of the gene shown on the right (CoV-2 and other CoVs: genes involved in the entry of these viruses, other categories: marker genes for specific cell types such as horizontal basal cells (HBC), and sustentacular cells (SUS)). Mean ± SD normalized counts in OSNs — Ace2: 8.6 ± 4.2, Tmprss2 117.3 ± 24.7, Ctsb: 38616.7 ± 1650.2, Ctsl: 1705 ± 87; same in WOM — Ace2: 254.7 ± 22.5, Tmprss2 2279 ± 219.6, Ctsb: 22380 ± 947, Ctsl: 4900 ± 90.5) **(B)** Log_2_-fold change (FC) of gene expression between OSNs and WOM as in Fig. 2A for three bulk RNA-sequencing datasets. MHV, mouse hepatitis virus. Left plot is same as Fig. 2A except for the addition of Ceacam1. **(C)** Gene expression for CoV-related genes including Ace2 and Tmprss2 as well as marker genes for olfactory and RE subtypes are shown normalized by their maximum expression across cell types. Ace2 and Tmprss2 are expressed in WOM respiratory and non-neuronal olfactory cell types, but not in OSNs. **(D)** UMAP representations of gene expression in the WOM dataset for CoV-2 related genes Ace2 and Tmprss2, as well as marker genes for each cell type. Each point represents an individual cell, and the color represents the normalized expression level for each gene (number of UMIs for a given gene divided by the total number of UMIs for each cell). **(E)** Fluorescent in situ hybridization of an identified dorsal sustentacular cell marker, Sult1c1 (in yellow), combined with immunostaining for the known dorsal OSN marker NQO1 (white). Note that Sult1c1 RNA fills the apical cytoplasm; given that sustentacular cells are ubiquitous in the epithelium, this is apparent as broad antisense signal for Sult1c in a pattern that is characteristic of the apical anatomy of sustentacular cells. Sult1c1 RNA is detected in sustentacular cells in the NQO1-positive dorsal olfactory epithelium. Nuclei were stained with DAPI (blue). Bar = 20 μm.

**Fig. S4.**
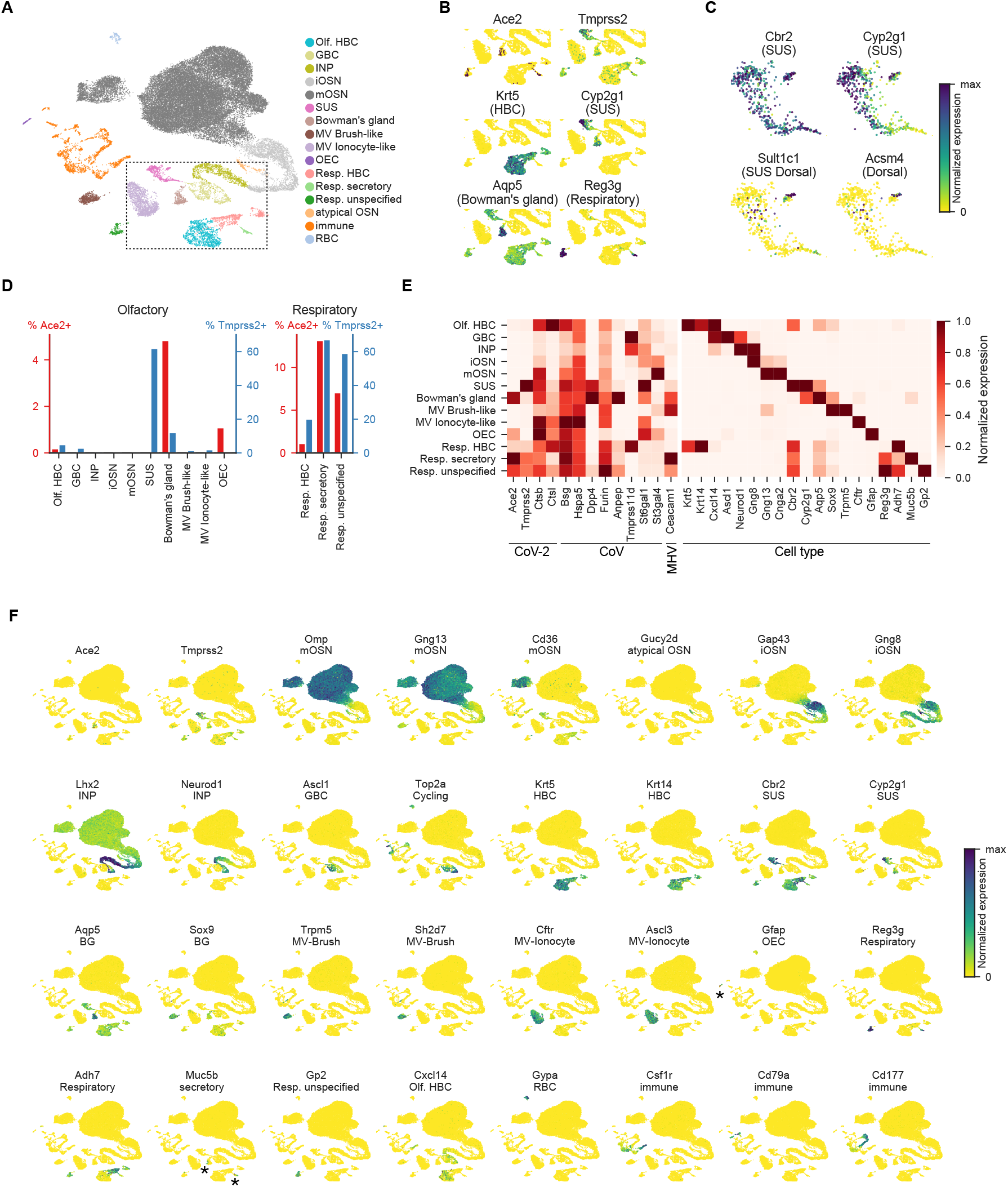
Related to Fig. 3. Analysis of a mouse WOM scSeq (10x chromium) dataset. **(A)** UMAP representation of single cell transcriptome data from WOM, colored by cell types (mOSN: mature OSN, iOSN: immature OSN, SUS: sustentacular cell, MV: microvillar cell, OEC: olfactory ensheathing cell, Resp.: respiratory, RBC: red blood cell). **(B)** UMAP representation of cell types shown in bounding box in A (plus the unspecified respiratory cluster, with expression of CoV-2 cell entry-related genes Ace2 and Tmprss2, as well as marker genes for HBCs, SUS cells, Bowman’s gland cells, and respiratory cells indicated. Each point represents an individual cell, and the color represents the normalized expression level for each gene (number UMIs for a given gene divided by the total number of UMIs for each cell). Only the cells in the area shown by black dashed box in **A** are plotted. **(C)** UMAP representation of gene expression in sustentacular cells, with pan-sustentacular and dorsal marker genes indicated. Note that only a small number of dorsal sustentacular cells were detected in this dataset. **(D)** Percent of cells expressing Ace2 and Tmprss2 in cell types identified. Ace2 is detected in HBC, Bowman’s gland,Olfactory ensheathing cells (OEC) and respiratory cells. **(E)** Gene expression for CoV-related genes including Ace2 and Tmprss2 as well as marker genes for olfactory and RE subtypes are shown normalized by their maximum expression across cell types. Ace2 and Tmprss2 are expressed in WOM respiratory and olfactory cell types, but not in OSNs. **(F)** CoV-2 related genes Ace2 and Tmprss2, as well as marker genes for cell types in Fig. 2C., in UMAP representation of WOM dataset with normalized expression. Gfap-positive OECs (olfactory ensheathing cells) and Muc5b-positive secretory cells are indicated by asterisks.

**Fig. S5.**
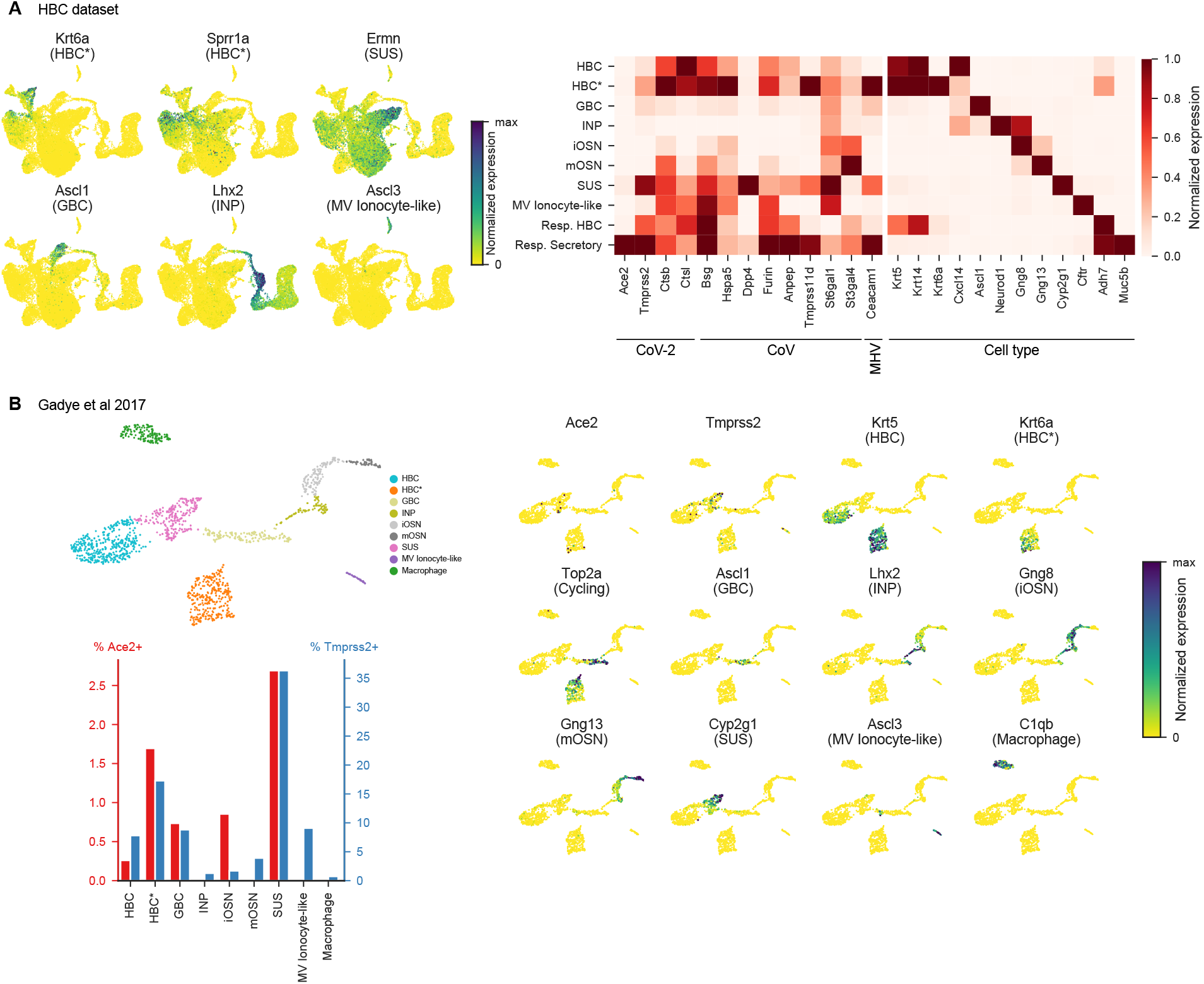
Related to Fig. 4. Analysis of a mouse Horizonal Basal Cell (HBC) lineage scSeq dataset, and analysis of publicly available HBC dataset. **(A)** (left) UMAP representation of cells belonging to a HBC lineage dataset (obtained after methimazole-mediated epithelial damage) with normalized expression (number of UMIs for a given gene divided by the total number of UMIs for each cell) for the indicated marker genes (GBC = globose basal cell, INP = intermediate neural precursor, MV = microvillar cell). (right) Gene expression levels for CoV-related genes including Ace2 and Tmprss2, as well as marker genes for olfactory and RE subtypes, normalized by their maximum expression across cell types (iOSN = immature olfactory sensory neuron, mOSN = mature olfactory neuron, SUS = sustentacular cell, Resp. = respiratory). Ace2 and Tmprss2 are expressed in WOM respiratory and olfactory cell types, but not in OSNs. HBC*= activated or cycling HBCs. **(B)** (top) UMAP representation of HBC lineage dataset from (*64*) colored by HBC lineage subtypes (top left) and (right) indicating normalized expression of identified marker genes. Each point represents an individual cell. (bottom left) Percent of cells expressing Ace2 and Tmprss2 in cell types identified in the HBC dataset. Ace2 is detected in sustentacular cells, horizontal basal cells, activated/cycling HBCs, globose basal cells, and iOSNs.

**Fig. S6.**
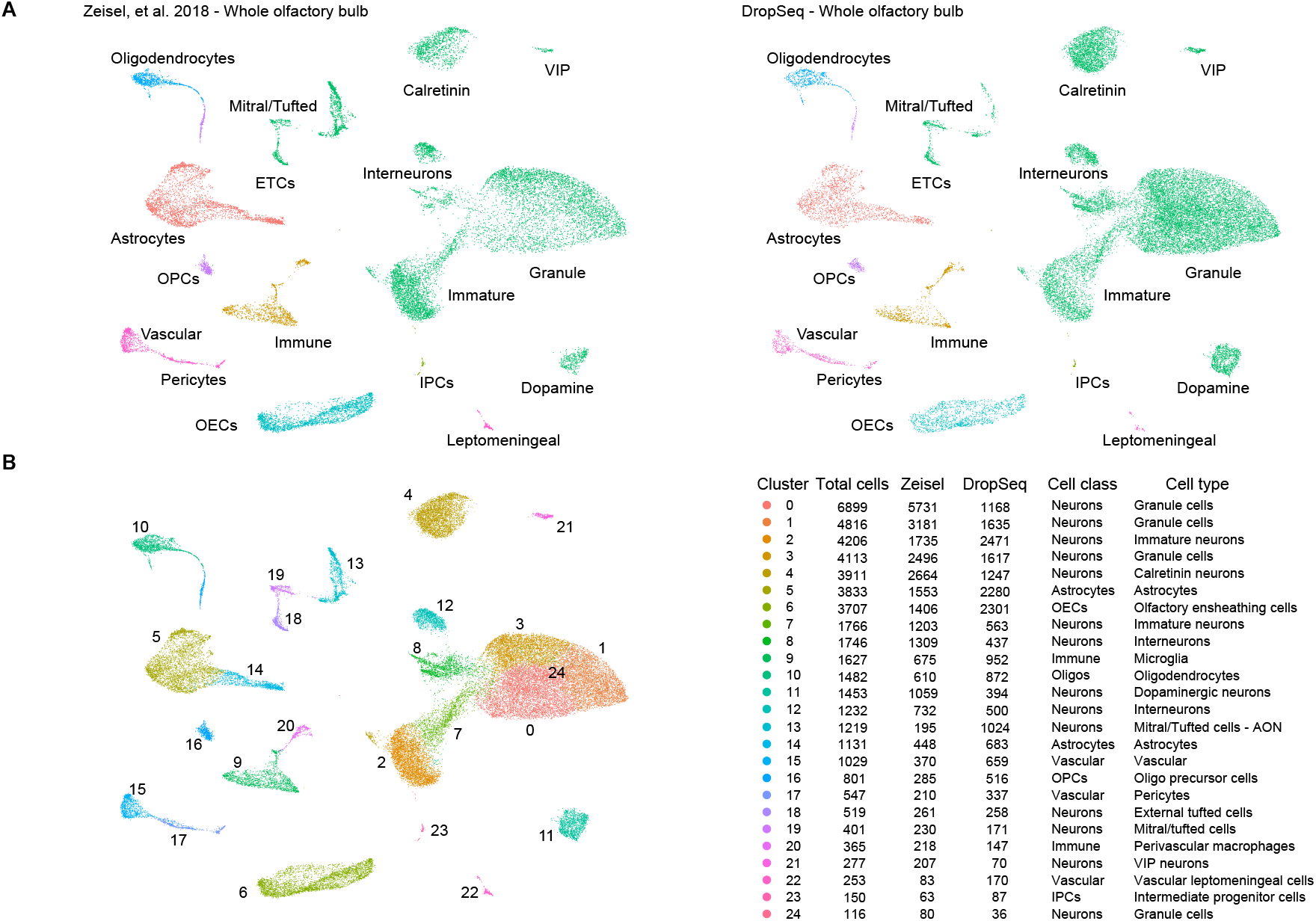
Related to Fig. 4. Whole OB scSeq data integration and clustering. **(A)** UMAP visualizations of olfactory bulb cells from Zeisel et al. (*42*)(left) and a new Drop-seq dataset (right). **(B)** Combined dataset UMAP visualization of clusters of OB cells (left), and table showing number of corresponding cells from each original dataset, cell classes and subtypes (right). VIP, vasoactive intestinal peptide positive neurons; ETCs, external tufted cells; OPCs, oligodendrocyte precursor cells; IPCs, intermediate precursor cells; OECs, olfactory ensheathing cells; AON, anterior olfactory nucleus.

**Fig. S7.**
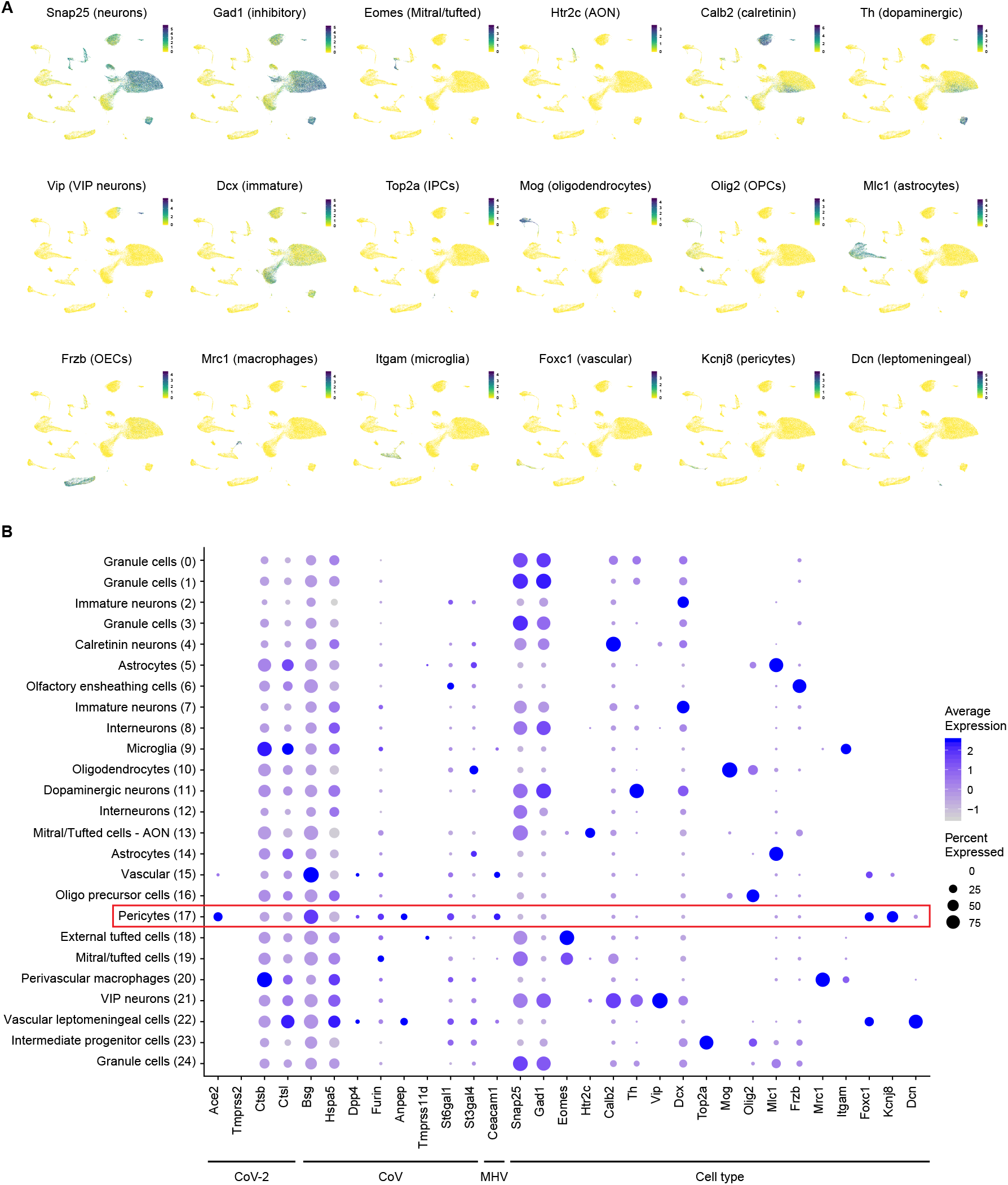
Related to Fig. 4. Expression of cell type markers in whole OB scSeq data. **(A)** UMAP visualisations showing expression of markers for neurons (Snap25), inhibitory neurons (Gad1), mitral and tufted cells (Eomes), excitatory neurons from the anterior olfactory nucleus (AON; Htr2c), glomerular layer calretinin positive neurons (Calb2), glomerular layer dopaminergic neurons (Th), vasoactive intestinal peptide (VIP) positive neurons (Vip), immature neurons (Dcx), intermediate progenitor cells (IPCs; Top2a), oligodendrocytes (Mog), oligodendrocyte precursor cells (OPCs; Olig2), astrocytes (Mlc1), olfactory ensheathing cells (OECs; Frzb), perivascular macrophages (Mrc1), microglia (Itgam), vascular cells (Foxc1), pericytes (Kcnj8) and vascular leptomeningeal cells (Dcn). **(B)** Dot plot showing expression of cell class and subtype markers alongside genes coding for coronavirus entry proteins in whole OB scSeq data. Circle size denotes the percentage of cells in each class expressing the gene; for plotting, minimum expression was set to 0.01% of cells per cluster. Color scale shows mean scaled expression level per cluster.

**Fig. S8.**
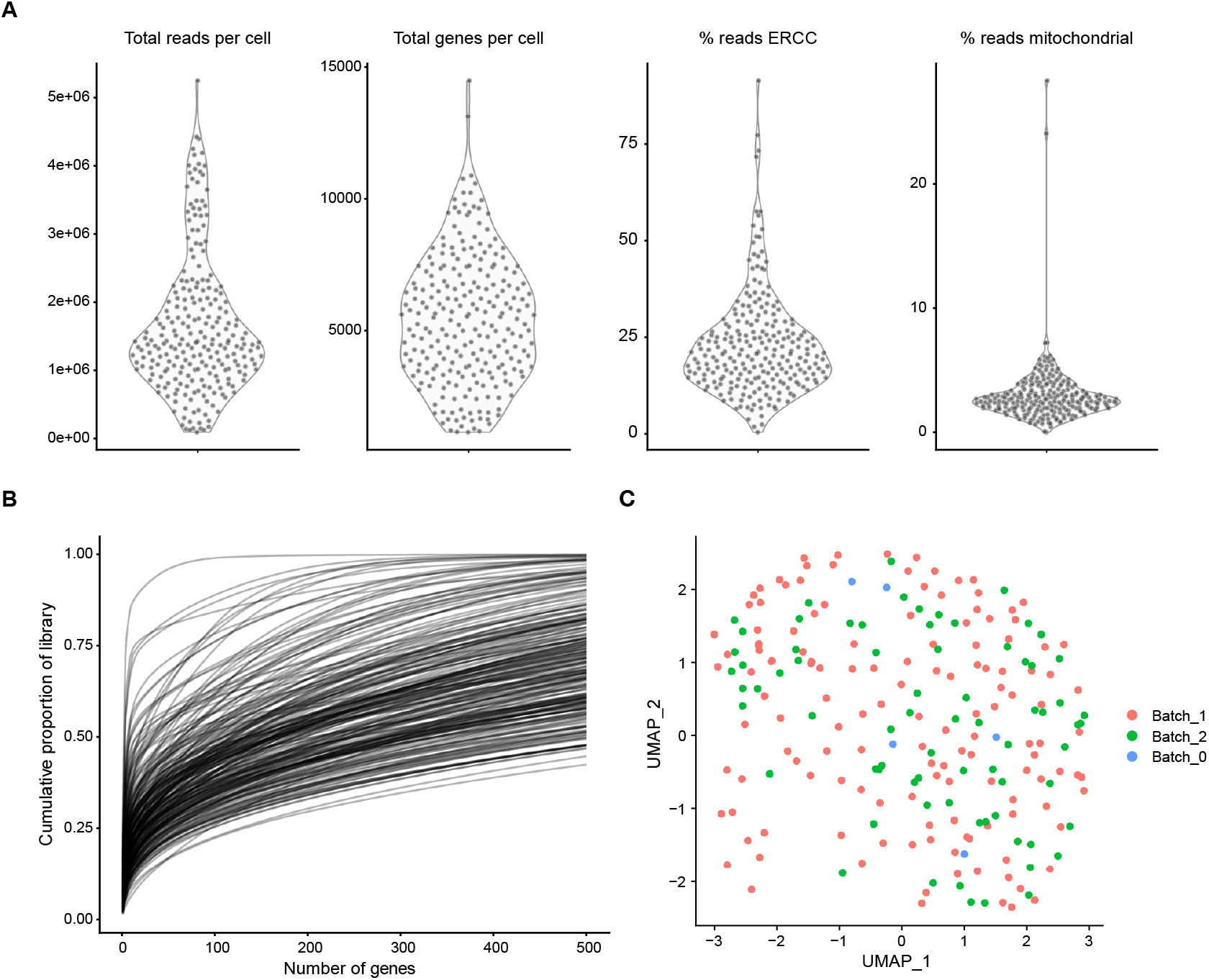
Related to Fig. 4. Sample quality control of manually sorted OB dopaminergic neurons. **(A)** Sample quality control metrics showing total number of reads per cell (left), total genes per cell (centre-left), percentage of reads per cell mapping to ERCC spike-ins (centre-right) and percentage of reads per cell mapping to mitochondrial genes (right). **(B)** Per cell library complexity depicted as the cumulative proportion of detected genes. **(C)** UMAP visualization of manually sorted DA neurons processed on three separate batches of library preparation and sequencing, showing efficient correction of technical batch effects with the scTransform function in Seurat (see Methods).

**Fig. S9.**
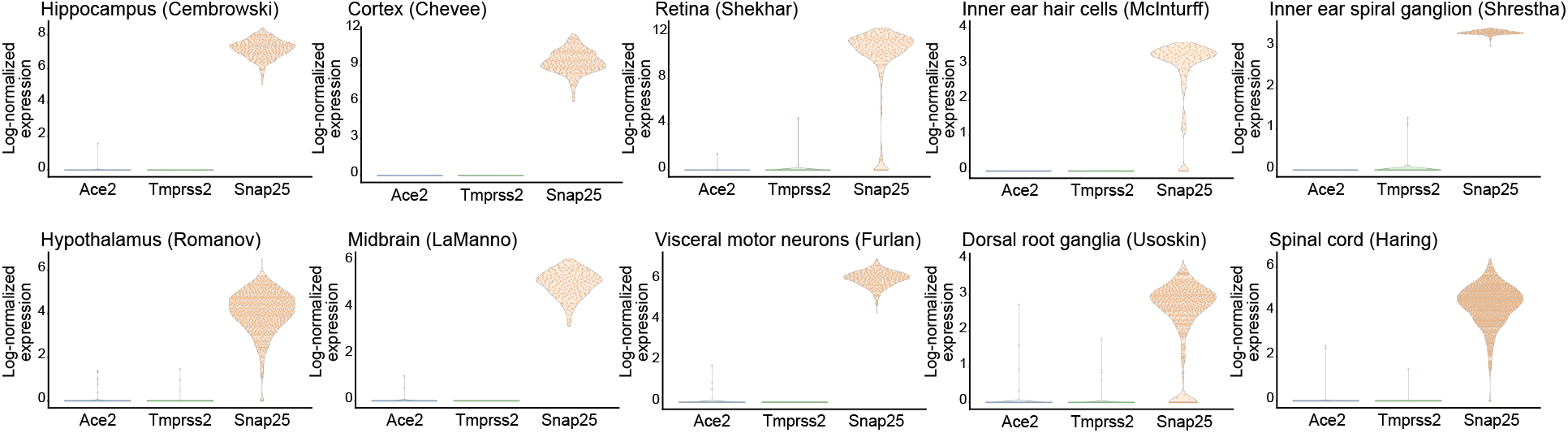
Related to Fig. 4. Expression of genes coding for CoV-2 entry-related proteins in neurons forming part of deeply sequenced scSeq datasets from various brain regions and sensory systems. Violin plots show expression of Ace2 and Tmprss2, alongside the neuronal marker Snap25 for reference. Expression values are log_2_-normalized counts of the number of transcripts for each gene. Each dot represents a single cell. Ace2 and Tprss2 are both only rarely detected in neurons.

